# Next-Generation Mapping of the ACINUS-Mediated Alternative Splicing Machinery and Its Regulation by O-glycosylation in *Arabidopsis*

**DOI:** 10.1101/2025.01.04.631329

**Authors:** Ruben Shrestha, Andres V Reyes, Shane Carey, Sumudu S Karunadasa, Wenxuan Zhai, Danbi Byun, Wen-Dar Lin, Jie Li, Kathrine Alerte, Hongchang Cui, Zhi-Yong Wang, Shou-Ling Xu

**Affiliations:** Department of Plant Biology, Carnegie Institution for Science, Stanford, CA, USA; Institute of Plant and Microbial Biology, Academia Sinica, 115 Taipei, Taiwan; Department of Biological Science, Florida State University, Tallahassee, Florida 32306, USA

## Abstract

Alternative splicing (AS) is a key mechanism of gene regulation, but the full repertoire of proteins involved and the regulatory mechanisms governing this process remain poorly understood. Using TurboID-based proximity labeling coupled with mass spectrometry (PL-MS), we comprehensively mapped the Arabidopsis AS machinery, focusing on the evolutionarily conserved splicing factor ACINUS, its paralog PININ, and the stable interactor SR45. We identified 298 high-confidence components, including both established and novel interactors, providing strong evidence that alternative splicing is coupled to transcription and that multiple RNA processing steps occur simultaneously in plants. Bioinformatic analysis reveals high redundancy, conserved mechanisms, and unique plant-specific features. Selected known and novel interactors were validated by AS readouts and phenotypic analysis, which also revealed a coordinated influence on splicing. Furthermore, a systematic evaluation of O-glycosylation double mutants revealed that SECRET AGENT (O-GlcNAc transferase) and SPINDLY (O-fucose transferase) modulate AS through both ACINUS-dependent and -independent pathways. Our results reveal the conserved as well as plant-specific AS regulatory network and highlight the global role of sugar modification in RNA processing.

## Introduction

Alternative splicing (AS) is a key mechanism of gene regulation in both animals and plants (1). During AS, the splicing machinery removes the introns by selecting from competing splice sites within the pre-mRNA, which is controlled by interactions between cis-acting elements and trans-acting factors, such as RNA-binding proteins (RBPs) and splicing factors. In a context-dependent manner, these factors either promote or inhibit splice site recognition and spliceosome assembly. AS generates multiple mRNA variants and protein isoforms from a single gene, affecting approximately 95% of genes in animals and 70% in plants (2–5).

The biological importance of AS has been demonstrated in several species. In flies, AS is essential for sex determination (6), while in humans it plays a critical role in tissue and organ development (7,8), with its dysregulation often associated with diseases such as cancer (9–11). In plants, AS is critical for establishing tissue identity, enhancing immunity, and responding to environmental changes and developmental signals (5,12–15).

Characterization of the proteins that control AS is essential for understanding how this complex system accurately distinguishes between multiple, often similar, splice sites to precisely remove the introns. The spliceosome, the molecular machinery responsible for splicing, is one of the most sophisticated megadalton complexes in eukaryotic cells (16). It orchestrates two transesterification reactions that result in lariat formation, intron removal, and exon ligation. The spliceosome consists of five small nuclear RNAs (snRNAs) that assemble with proteins to form small nuclear ribonucleoproteins (snRNPs), along with hundreds of additional proteins, including non-snRNP splicing factors, cofactors and regulatory proteins (17). Despite extensive research ranging from proteomic analyses using immunoprecipitations to systematic gene silencing/knockout using RNAi or CRISPR-Cas approaches, the complete composition and dynamic behavior of the spliceosome remain incompletely elucidated (18–20).

While the core components of the splicing machinery, such as snRNPs and certain non-snRNP proteins, are highly conserved among eukaryotes, AS exhibits both conserved (21) and species-specific features (22). A key difference is intron length: Arabidopsis introns are significantly shorter (median length of 100 nucleotides) (23) than human introns (median length of 1.5-1.7 kb) (20,24). This disparity likely reflects differences in the mechanisms of snRNP and SR proteins binding to pre-mRNA. In Arabidopsis, it has been suggested that the splicing machinery predominantly uses intron definition, where the splicesome first recognizes and assembles across introns. In contrast, exon definition, where the splicesome first recognizes and assembles across exons - is more common in humans (20). These mechanistic differences correlate with different patterns of AS: in plants, intron retention is the most common form of AS, whereas in humans, exon skipping is the dominant AS event (5,25,26). A comprehensive understanding of the proteins involved in plant AS will provide crucial insights into the mechanisms and evolution of AS.

Post-translational modifications (PTMs), such as phosphorylation, have been shown to play a critical role in the regulation of AS (27,28). Inhibitor treatments of O-GlcNAc transferase OGT and O-GlcNAcase OGA cause a global change in retained intron levels, although the underlying mechanism remains elusive (29). In addition to phosphorylation, our results indicate that several splicing factors are modified by O-GlcNAcylation and O-fucosylation (30–32). qPCR data reveal AS defects in a few genes in single mutants of the O-GlcNAc transferase SECRET AGENT (SEC) and the O-fucose transferase SPINDLY (SPY) single mutants (13), suggesting that O-glycosylation plays a role in AS regulation. However, a comprehensive evaluation of the global effects of SEC and SPY on AS, including the analysis of double mutants, is still lacking. Such studies are needed to elucidate the role of O-GlcNAcylation and O-fucosylation in AS regulation.

Previously, we identified the evolutionarily conserved splicing factor ACINUS and its paralog PININ as key regulators of transcription, AS, and developmental transitions (13). In this study, we used TurboID, an engineered high-activity biotin ligase (33,34), to comprehensively map ACINUS-mediated AS complexes. By constructing a high-confidence network centered on three baits-ACINUS, its paralog PININ, and the stable interactor SR45(a homolog of RNPS1) (13,35–38)-we characterized the composition of the AS machinery and identified both established and novel interactors, conserved mechanism as well as plant-specific features. We validated the phenotypic readouts of several interactors, using either molecular readouts (alternative splicing defects) or morphological phenotypes. We further uncovered the roles of O-GlcNAcylation and O-fucosylation in regulating AS and transcription in conditional *spy sec* double mutants, suggesting both ACINUS-dependent and -independent mechanisms. These findings provide a systematic framework for understanding the AS machinery in plants and uncovering how AS is regulated, paving the way for further exploration of AS mechanisms in plant systems.

## Results

### Improved sequencing reveals widespread alternative splicing defects in *acinus pinin* mutants

Our previous single-read (100 bp) sequencing with an average coverage of 22.4 million reads identified approximately 289 altered retained intron events in *acinus pinin* mutants, along with a few other AS events, using RackJ (Table S1) (13). To improve AS detection, we performed new paired-end (150bp) non-stranded sequencing for both WT and *acinus pinin* mutants (n=4), achieving an average of 57.2 million reads per replicate (Table S1). Higher coverage and paired-end reads will improve the detection of exon-exon and exon-intron junctions, thus improving the analysis of AS (39).

We also refined the detection of retained introns by integrating the intron read <u>d</u>epth <u>ratio</u> (IDratio) into RackJ (Fig. S1A). IDratio quantifies cDNA with retained introns relative to the total cDNA for a given gene, based on the read depths of introns and flanking exons. This metric provides a more intuitive measure of intron retention than our previous intron coverage ratio (ICratio); for example, an IDratio of 10% indicates that 10% of a gene’s transcripts contain retained introns. In addition, we applied more stringent thresholds (Fig.S1A-C), including a mean IDratio of at least 15% in at least one group (WT or *acinus pinin*) for statistical analysis.

Our analysis revealed extensive AS defects in *acinus pinin* mutants compared to wild type using a 1.5-fold cutoff (Table 1), with 1,106 altered retained intron events, along with 206 exon skipping events, 371 alternative donor events, 498 alternative acceptor events, and 265 combined alternative donor and acceptor, in the double mutants (Table 1). These results highlight the critical role of ACINUS and PININ in AS regulation.

**Table 1:**
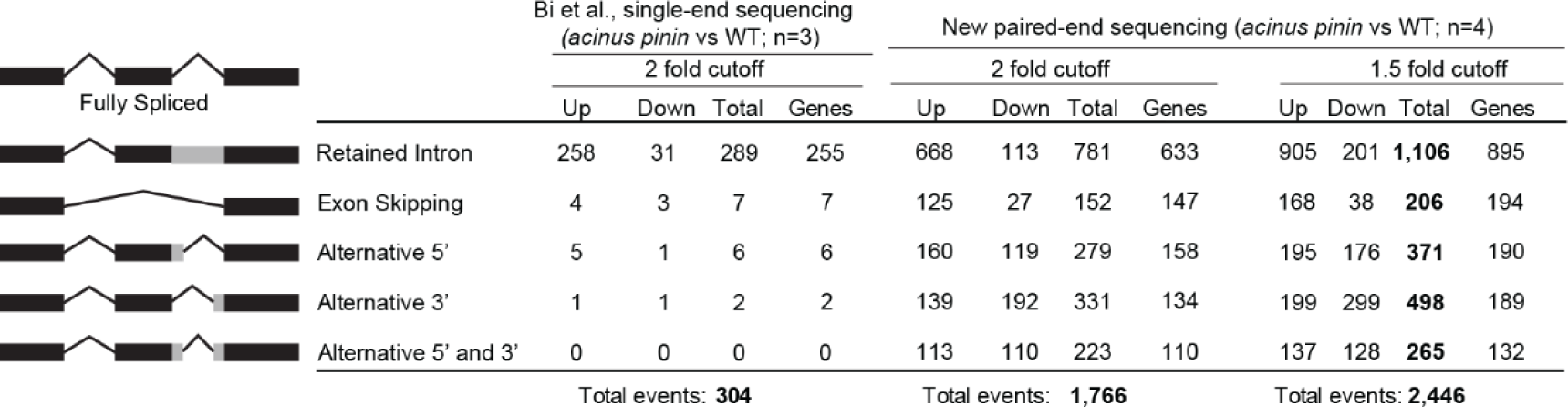
Improved sequencing reveals widespread alternative splicing defects in *acinus pinin* mutants. . Paired-end (150bp) non-stranded sequencing of WT and *acinus pinin* mutants (n=4), with an average of 57.2 million reads per replicate, significantly improves the detection of alternative splicing events. More stringent thresholds for intron retention analysis were applied to the new sequencing data, and the number of different AS events altered in *acinus pinin* mutants using 2-fold and 1.5-fold cutoffs are presented.

|  |  | Bi et al., single-end sequencing<br>(*acinus pinin* vs WT; n=3) |  |  |  | New paired-end sequencing (*acinus pinin* vs WT; n=4) |  |  |  |  |  |  |  |
| --- | --- | --- | --- | --- | --- | --- | --- | --- | --- | --- | --- | --- | --- |
|  |  | 2 fold cutoff |  |  |  | 2 fold cutoff |  |  |  | 1.5 fold cutoff |  |  |  |
|  |  | Up | Down | Total | Genes | Up | Down | Total | Genes | Up | Down | Total | Genes |
|  | Fully Spliced |  |  |  |  |  |  |  |  |  |  |  |  |
|  | Retained Intron | 258 | 31 | 289 | 255 | 668 | 113 | 781 | 633 | 905 | 201 | **1,106** | 895 |
|  | Exon Skipping | 4 | 3 | 7 | 7 | 125 | 27 | 152 | 147 | 168 | 38 | **206** | 194 |
|  | Alternative 5' | 5 | 1 | 6 | 6 | 160 | 119 | 279 | 158 | 195 | 176 | **371** | 190 |
|  | Alternative 3' | 1 | 1 | 2 | 2 | 139 | 192 | 331 | 134 | 199 | 299 | **498** | 189 |
|  | Alternative 5' and 3' | 0 | 0 | 0 | 0 | 113 | 110 | 223 | 110 | 137 | 128 | **265** | 132 |
|  |  | Total events: **304** |  |  |  | Total events: **1,766** |  |  |  | Total events: **2,446** |  |  |  |

### Profile ACINUS proxiome using TurboID-based PL-MS and functional analysis

To gain a reliable and comprehensive understanding of the plant alternative splicing machinery, and the regulatory roles of ACINUS and PININ, we performed TurboID-based proximity labeling proteomic profiling (Fig. 1A) to map their interactomes. We fused ACINUS, PININ, and SR45 to TurboID-VENUS and expressed them under their native promoters in *acinus-2 pinin* double mutant or *sr45-1* mutant backgrounds (Fig. 1B). To ensure specificity, we included two controls: nuclear-localized TurboID fused to YFP (TD-YFP) as a spatial reference control and ACINUSΔRSB-TD, a variant lacking the 592-633 RSB domain (40), both expressed in wild-type plants (Fig. 1B).

**Figure 1.**
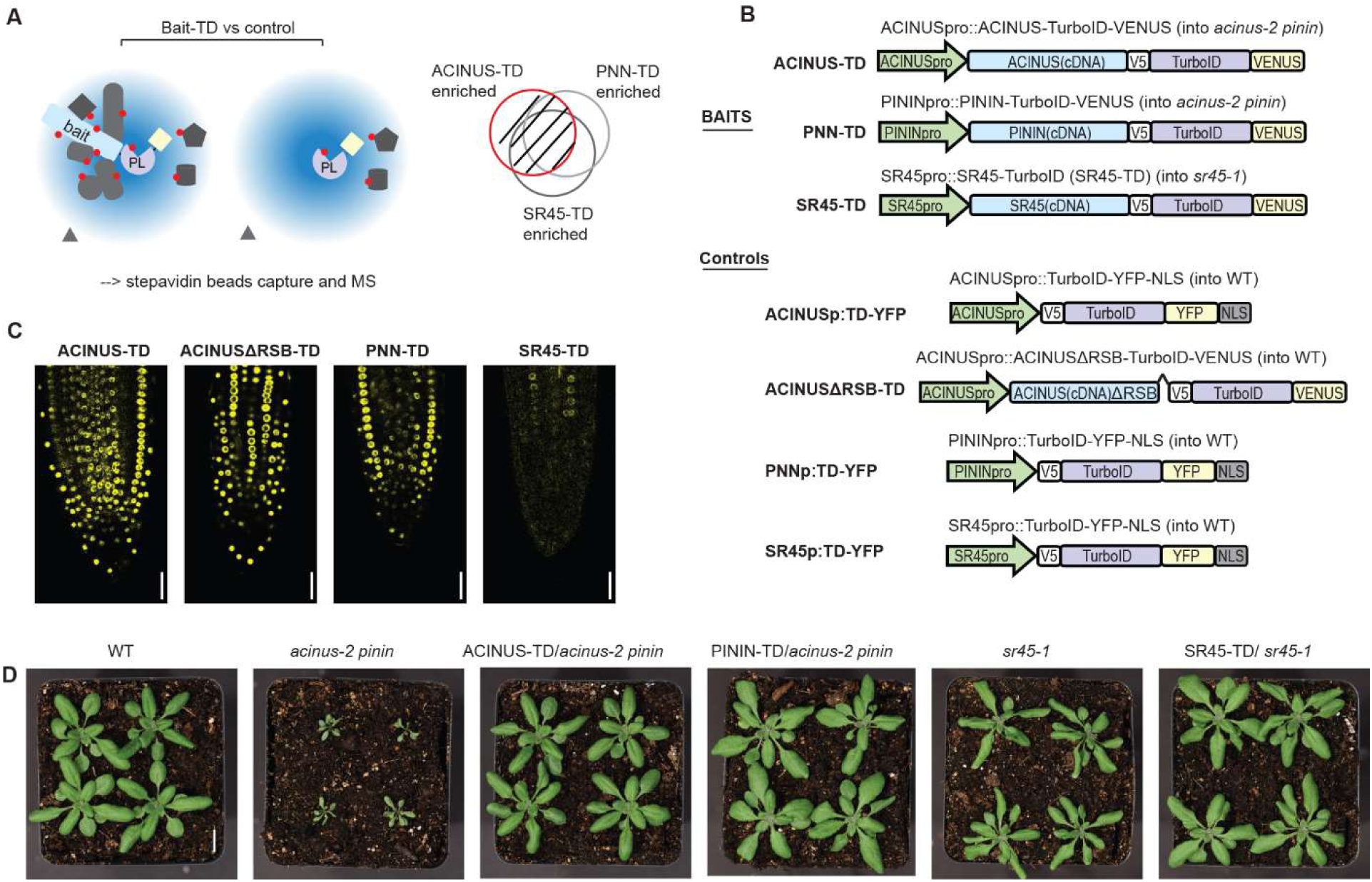
Schematic of PL-MS for ACINUS proxiome mapping and functional analysis. (A) Schematic of TurboID-based interactome mapping. Three bait proteins - ACINUS, PININ, SR45 - are fused to TurboID (TD) and VENUS, while the control is TD-YFP, expressed under native promoter. Proteins within the labeling radius (blue cloud) are labeled with biotin (red dot), then enriched with streptavidin beads followed by mass spectrometric analysis. After comparison with the control, the enriched protein list from each bait is overlapped and filtered to generate a highly reliable interactome list. (B) Construct information of the bait and corresponding control. All constructs contain V5 and VENUS or YFP tags. Controls contain a nuclear localization signal (NLS). (C) Confocal microscopy shows that the ACINUS-TD, ACINUSΔRSB-TD, PININ-TD, and SR45-TD fusion proteins localize to the nucleus. (D) ACINUS-TD, PNN-TD and SR45-TD fusion proteins rescued the corresponding mutants to wild-type levels.

Confocal and spinning disk microscopy confirmed the nuclear localization of all three proteins, with ACINUS being the most abundant and SR45 the least abundant (Fig. 1C, and Fig. S2A). To assess the physiological functionality of the full-length fusion proteins, we evaluated leaf shape, rosette size, and flowering time in these lines. The *acinus pinin* and *sr45* mutants exhibit pleiotropic phenotypes, including narrow, twisted rosette leaves with serrated margins, small rosettes, and delayed flowering, with the *acinus pinin* mutants exhibiting a more severe phenotype. As expected, the full-length fusion constructs completely rescued these mutant phenotypes to wild-type levels, confirming that the fusion proteins are functional in planta (Fig. 1D, Fig. S2B-C).

### Comprehensive high-confidence PL-MS mapping of the ACINUS-proxiome

We performed PL-MS on the ACINUS-TD, PNN-TD, and SR45-TD lines using protein-level enrichment for capturing intact biotinylated proteins (41). All three lines, along with their control lines, were treated with 50 µM biotin for 1 hour, with three biological replicates each. Biotinylated proteins were enriched using streptavidin beads, measured by LC/MS/MS, and quantified using label-free quantification (LFQ) (Fig. 2A). Using a threshold of S0=2 and an FDR of 0.05 in Perseus, we identified 395, 324, and 255 proteins enriched by ACINUS-TD, PININ-TD, and SR45-TD, respectively, with known interactors (in red font) and novel interactors (in blue font) selectively labeled (Fig. 2B-D, Fig. S3A). The higher number of proteins enriched by ACINUS-TD is likely due to its higher expression, resulting in a higher signal-to-noise ratio, as indicated by the enrichment ratio difference shown on the x-axis, compared to PININ-TD and SR45-TD (Fig. 2B-D). Therefore, we prioritized ACINUS for data integration.

**Figure 2.**
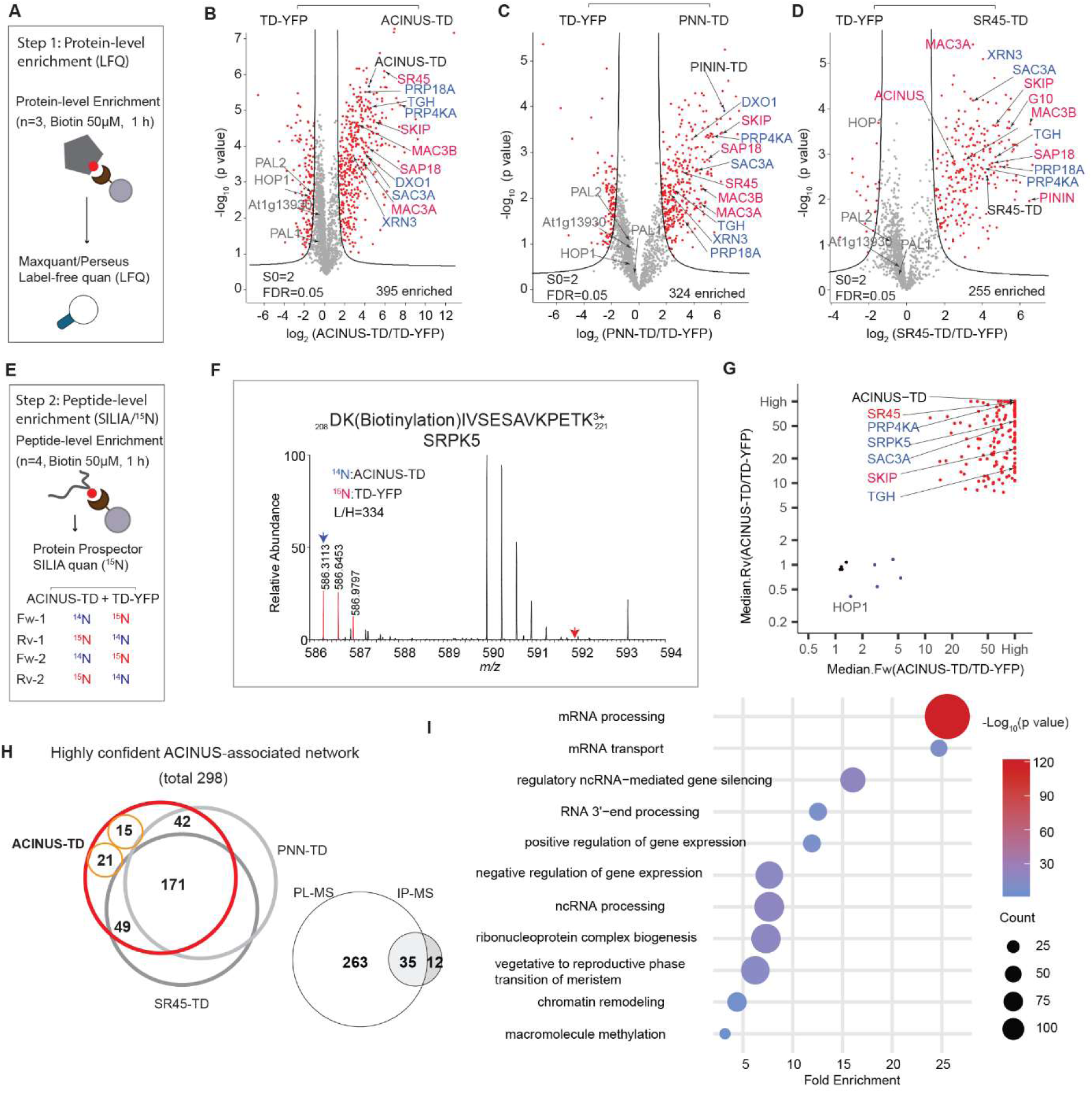
Comprehensive high-confidence PL-MS mapping of the ACINUS proxiome. (**A**) Schematic of the workflow for mapping the ACINUS proxiome at the protein level. Baits and controls were treated with biotin (n=3 biological replicates), followed by protein-level enrichment to capture intact biotinylated proteins and label-free quantification (LFQ). (**B**-**D**) Volcano plots of ACINUS-TD, PNN-TD, and SR45-TD. Proteins significantly enriched by either the bait (right) or control (left) are highlighted in red. Bait proteins, selected known and novel interactors are labeled in black, red, and blue, respectively. Certain standby nuclear proteins biotinylated by both Turbo-YFP control and bait are labeled in gray. (E) Schematic of the workflow for ACINUS peptide-level proxiome mapping. After SILIA labeling, protein extraction, and tryptic digestion, biotinylated peptides were enriched using streptavidin beads, followed by ^15^N quantification. (F) Example of SILIA-based quantification. The quantitative difference of a biotinylated peptide from SRPK5 in the mixed sample (^14^N: ACINUS-TD; ^15^N: TD-YFP) is shown. (G) Quantification of ACINUS-TD after peptide-level enrichment. Bait-enriched proteins are highlighted in red, biotinylated standby nuclear proteins and endogenous biotinylated proteins are highlighted in blue, and unmodified proteins showing equal mixing are highlighted in black. Bait proteins and selected known and novel interactors are labeled in black, red, and blue, respectively. (H) Plot of a high-confidence interactome was generated by extensive filtering and comparison of TurboID-based PL-MS with IP-MS. Numbers represent proteins enriched by ACINUS-TD that overlap with those enriched by both PININ-TD and SR45-TD (n=171 proteins, including the three baits), overlap exclusively with PNN-TD (n=42) or SR45-TD (n=49), enrichment by ACINUS-TD at both protein and peptide level (n=15), and enrichment at the peptide level in at least 3 out of 4 replicates (n=21). The filtered high-confidence of ACINUS-TD proxiome (298 proteins) overlaps with ACINUS IP-MS, with 35 shared proteins. (I) Go analysis of the ACINUS proxiome shows significant enrichment in mRNA processing and other processes.

Next, we used a complementary approach to directly capture and quantify biotinylated peptides enriched by ACINUS-TD using ^15^N stable isotope labeling in Arabidopsis (SILIA) quantitative mass spectrometry (42). ACINUS-TD and TD-YFP lines were metabolically labeled with ^14^N and ^15^N, respectively, and treated with 50 µM biotin for 1 h, followed by biotinylated peptide enrichment and ^15^N quantification over four biological replicates (Fig. 2E-2G, Fig. S3B). To ensure high confidence, the MS/MS peak lists were first filtered using the biotin signature ion (derivative ion at m/z 310.16 (^14^N) or 311.16 (^15^N) due to ammonia loss from the immonium ion of biotinylated lysine (ImKBio) (Fig. S3C-D), prior to subsequent search and quantification for biotinylated peptides. Biotinylated peptides identified by both ^14^N and ^15^N labeling from ACINUS-TD enriched samples were considered high confidence candidates. This analysis identified 156 significantly enriched proteins in the ACINUS-TD samples that were quantified in at least three out of four replicates (Fig. 2G). While the peptide-level enrichment showed substantial overlap with the protein-level enrichment (Table S2), it also uncovered unique biotinylated proteins, such as SERINE/ARGININE PROTEIN KINASE 5(SRPK5), providing complementary insights (Fig. 2F).

To effectively filter and integrate both datasets, we implemented a stringent selection system to generate a confidence interactome list (see Methods). We prioritized ACINUS-enriched proteins based on their overlap with those enriched by both PININ-TD and SR45-TD (n=171 proteins, including the three baits), exclusive overlap with PNN-TD (n=42) or SR45-TD (n=49), enrichment by ACINUS-TD at both protein and peptide level (n=15), and enrichment at the peptide level in at least 3 out of 4 replicates (n=21) (Fig. 2H). These rigorous filtering steps result in a total network of 298 proteins.

Notably, this high-confidence list includes 35 of the 47 proteins previously identified in our IP-MS using ACINUS as bait (Fig. 2H). Interestingly, 5 of the 12 missing proteins were significantly enriched by ACINUS-TD (Table S2) but were omitted due to our stringent criteria. Overall, this integrated TurboID dataset significantly extends the known ACINUS network and demonstrates exceptional sensitivity.

### Interdependence of AS, transcription, and other mRNA processing steps

To better understand the relationships between the identified interactors, we performed Gene Ontology (GO) term analysis (43) and used Cytoscape (44) to construct a comprehensive network, organizing proteins based on their associations and functional groups (Fig. S4). GO analysis shows that the ACINUS proxiome is significantly enriched for proteins involved in mRNA processing (Fig. 2I), including numerous homologs to known spliceosome components. We identified splicing factors with established or putative roles in splicing spanning multiple stages of splicesome assembly and catalysis (A, B and B^act^, and C complex), as well as key members of the NTC/PRP19 complex. In addition, our dataset captured core and accessory components of the spliceosome U1, U2, and U4/U6.U5 snRNP subunits, as well as numerous RS and hnRNPs proteins, underscoring the broad coverage of critical splicing regulators (Fig. 3, Fig. S4). Our ACINUS proxiome analysis provides strong evidence for the coupling between AS and transcription. Go analysis supports this association, showing significant enrichment in categories related to gene expression regulation, chromatin remodeling, and proteins associated with the vegetative to reproductive phase transition of the meristem (Fig. 2I). We identified a large number of key proteins involved in transcription and chromatin remodeling (Fig. 3), including RNA polymerase II elongation factors (TFIIS, TFIIE and TFIIEα) and global transcription factors (GTA2, GTB1, GTE4). In addition, we detected essential components of the polymerase-associated factor PAF1 complex (ELF5, ELF7, ELF8, VIP5, PHP) (45), chromatin remodelers (CHR5, CHR28, SWC4, SPT2) and RNA polymerase II C-terminal domain phosphatases (CPL1, CPL3). Our dataset also highlights the FAcilitates Chromatin Transcription (FACT) complex (HMG and SPT16), the histone H3-K4 methylation regulator (SDG2) (46) and the RDR2-independent DNA methylation proteins (NERD). These findings reveal an extensive and intricate network of molecular interactions linking transcription and AS, highlighting their coordinated roles in gene regulation.

**Figure 3.**
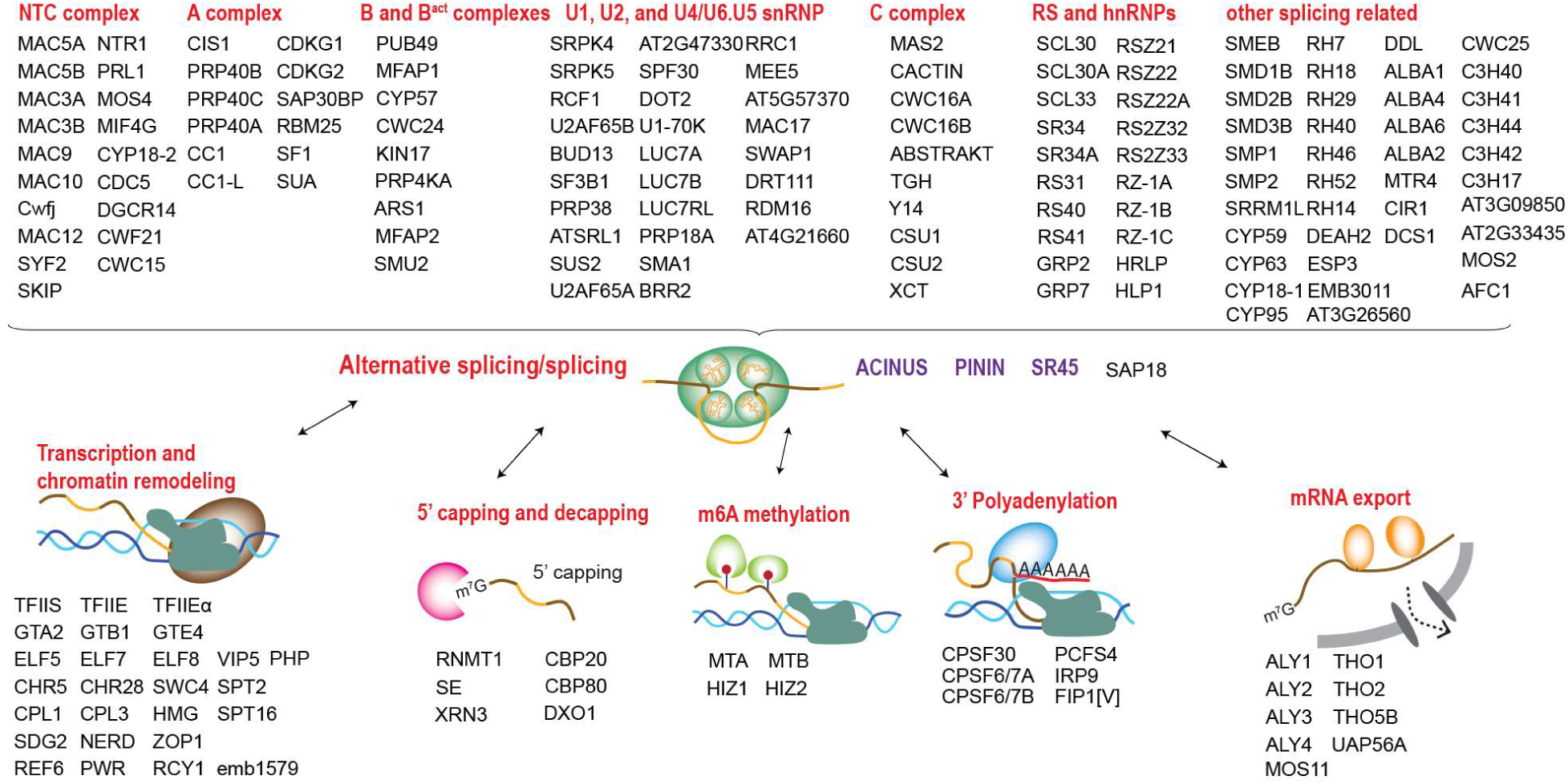
Interdependence of alternative splicing, transcription, and other mRNA processing steps. Enriched proteins in the ACINUS proxiome are classified based on the predicted functions, highlighting key factors with known or putative roles involved in splicing and alternative splicing, transcription and chromatin remodeling, 5’ capping and decapping, m6A methylation, 3’ polyadenylation, and mRNA export.

To investigate the relationship between AS and other RNA processing steps, we analyzed the proxiome and identified key proteins involved in 5’ capping and decapping, m6A methylation, 3’ polyadenylation, and mRNA export (Fig. 3). For 5’ capping, we detected the capping enzyme RNMT1, the capping binding protein complex (CBP20 and CBP80), the associated protein SE, and the decapping proteins XRN3 and DXO1, the latter known to activate RNMT1 for m7G capping (47). For m6A methylation, we identified the mRNA adenosine methylases MTA and MTB and the m6A writer complex HIZ1 and HIZ2. For 3’ polyadenylation, we identified CPSF30, CPSF6/7A, CPSF6/7B, PCFS4, IRP9 and FIP1[V]. In addition, we identified several components of the THO-TREX-1 complex that are essential for mRNA transport, including ALY1-4, THO1, THO2, THO5B, UAP56A, and MOS11. These findings reveal a tightly interconnected network of RNA processing events that are likely to ensure mRNA maturation and export.

### ACINUS-proxiome reveals high redundancy, conserved mechanisms, and unique plant-specific features

To investigate the features of the ACINUS proxiome, we analyzed the network using Interpro, which classifies proteins into families and predicts functional domains (48). This analysis identified features associated with AS and revealed extensive redundancy within the network. Notable domains include RNA binding domain (RBD), helicases, zinc finger motif (Znf_CCHC, Znf_CCCH, Znf_RING/FYVE/PHD), WD40 repeats, G-patch domain, WW domain, SURP motif, and K homology domain (KH) (Fig. 4A, Table S2). Several of these motifs have been shown to contribute to AS through different mechanisms: WD repeats, WW domains, and SURP motif facilitate protein-protein interactions; G-patch domain regulates RNA helicases; and KH domain and zinc finger motif mediate RNA or DNA binding. The presence of these features in animal counterparts (49) underscores the evolutionary conservation of AS mechanism between plants and animals. Notably, 21 helicases were detected, indicating the high redundancy within the network.

**Figure 4.**
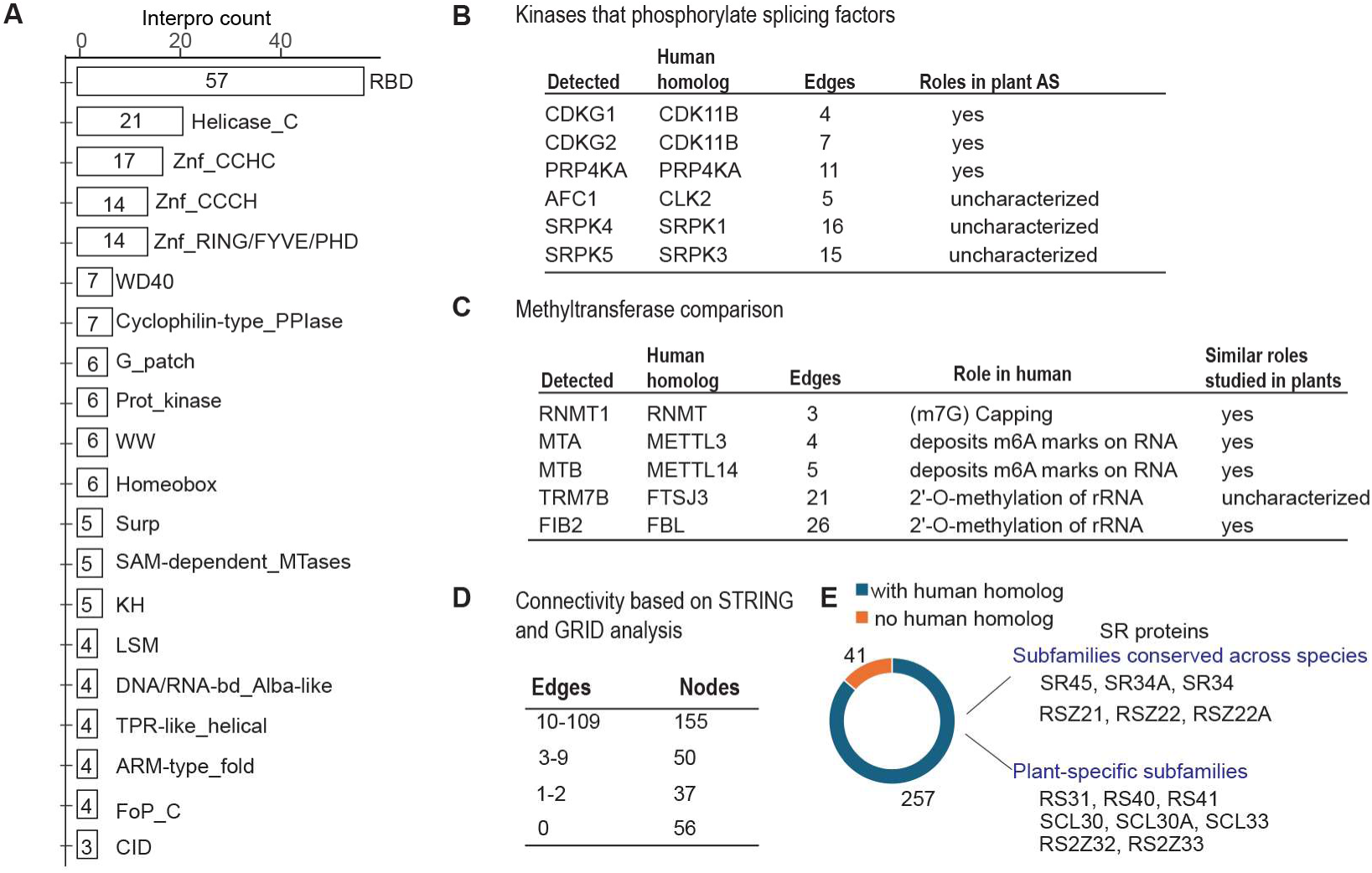
The ACINUS proxiome reveals high redundancy, conserved mechanisms, and unique plant-specific features. (A) Interpro analysis identifies features associated with AS and highlights extensive redundancy within the network. Numbers within the boxes indicate the number of proteins detected with specific domains, with domain names labeled on the right. (**B**-**C**) Kinases and methyltransferases detected in the ACINUS proxiome suggest conserved regulatory mechanisms in AS. (**D**) Connectivity summary of the ACINUS proxiome based on STRING and GRID analysis. (**E**) Blast analysis identified 257 proteins with human homologs and 41 without. Among 14 identified SR proteins, 6 belong to subfamilies conserved across species, while 8 are plant-specific.

We identified six kinases (Fig. 4B), all of which have human homologs involved in the regulation of AS (50). These include CLK kinases (CDKG1 and CDKG2), the splicing kinase PRP4KA, the LAMMER kinase family AFC1, and group II members of the SRPK family (SRPK4, SRPK5), suggesting a conserved regulatory mechanism between plants and animals. While CDKG1, CDKG2, and PRP4KA have established roles in AS regulation in plants, the functions of AFC1, SRPK4, SRPK5, and the specific substrates of all six kinases remain to be determined (51–54).

The identification of five methyltransferases further supports a conserved regulatory mechanism across species (Fig. 4C). Among these, RNMT1 (47), MTA, MTB (55), and FIB2 (56) have been shown in plants to have analogous functions to their human homologs in methylating mRNA or rRNA in plants, whereas TRM7B remains to be characterized. Given their association with the ACINUS proxiome, we suggest that methyltransferases, together with the identified kinases, play a similar role in AS regulation as their animal counterparts, and they are working on site of the complexes.

Cytoscape, InterPro and Blast analyses further revealed unique features of the ACINUS-proxiome. Cytoscape analysis revealed extensive connectivity within the network, with 155 proteins having 10-109 edges and 50 proteins having 3-9 edges. SF3B1 had the highest number of interactions with 109 edges (Fig. 4D, Fig. S4). In addition, we identified 93 proteins with much less connectivity, including 37 proteins with minimal prior connections (1-2 edges) and 56 proteins with no prior connections to the network (0 edges), suggesting a potential role in mRNA processing similar to ACINUS.

Blast analysis identified 255 proteins with human homologs, while 41 lacked homologs, including 12 without InterPro domains and 9 with plant-specific domains, highlighting adaptations unique to plant AS regulation (Fig. 4E, Table S2). Among those with human homologs, 14 were SR proteins, some of which belonged to conserved eukaryotic subfamilies such as the SR (SR45, SR34A and SR34) and RSZ (RSZ21, RSZ22, RSZ22A) subfamilies. In contrast, several SR proteins were plant-specific, including the RS (RS31, RS40, RS41), SCL (SCL30, SCL30A, SCL33), and RS2Z (RS2Z32 and RS2Z33) subfamilies (57), highlighting their specialized roles in plant AS regulation (Fig. 3 and 4E).

### The RSB domain is critical for ACINUS function as a hub protein

Previous studies have shown that the RSB domain interacts with RNPS1 (SR45 homolog) and SAP18 (40). To investigate the role of the RSB domain in the protein interaction network, we generated ACINUS-ΔRSB-TD transgenic lines with a deletion of the RSB domain 592-633 in the WT background. The truncated fusion protein was expressed at levels comparable to the full-length ACINUS-TD and localized to the nucleus (Fig. 1C) and did not cause any visible phenotypic changes.

We then compared the TurboID results of ACINUS-ΔRSB-TD to ACINUS-TD using protein-level enrichment. Surprisingly, we found that nearly 75% of the interactors showed either reduced or lost interactions (Fig. 5A-B, Fig. S5A). These affected interactors included SR45 and SAP18 and many other splicing factors. In addition, proteins involved in transcription and chromatin remodeling, and other RNA processing steps were significantly impacted (Fig. 5A-B, Fig. S4B, Table S2). This substantial loss of interactors underscores the critical role of the RSB domain for ACINUS as a network hub (Fig. 2G).

**Figure 5.**
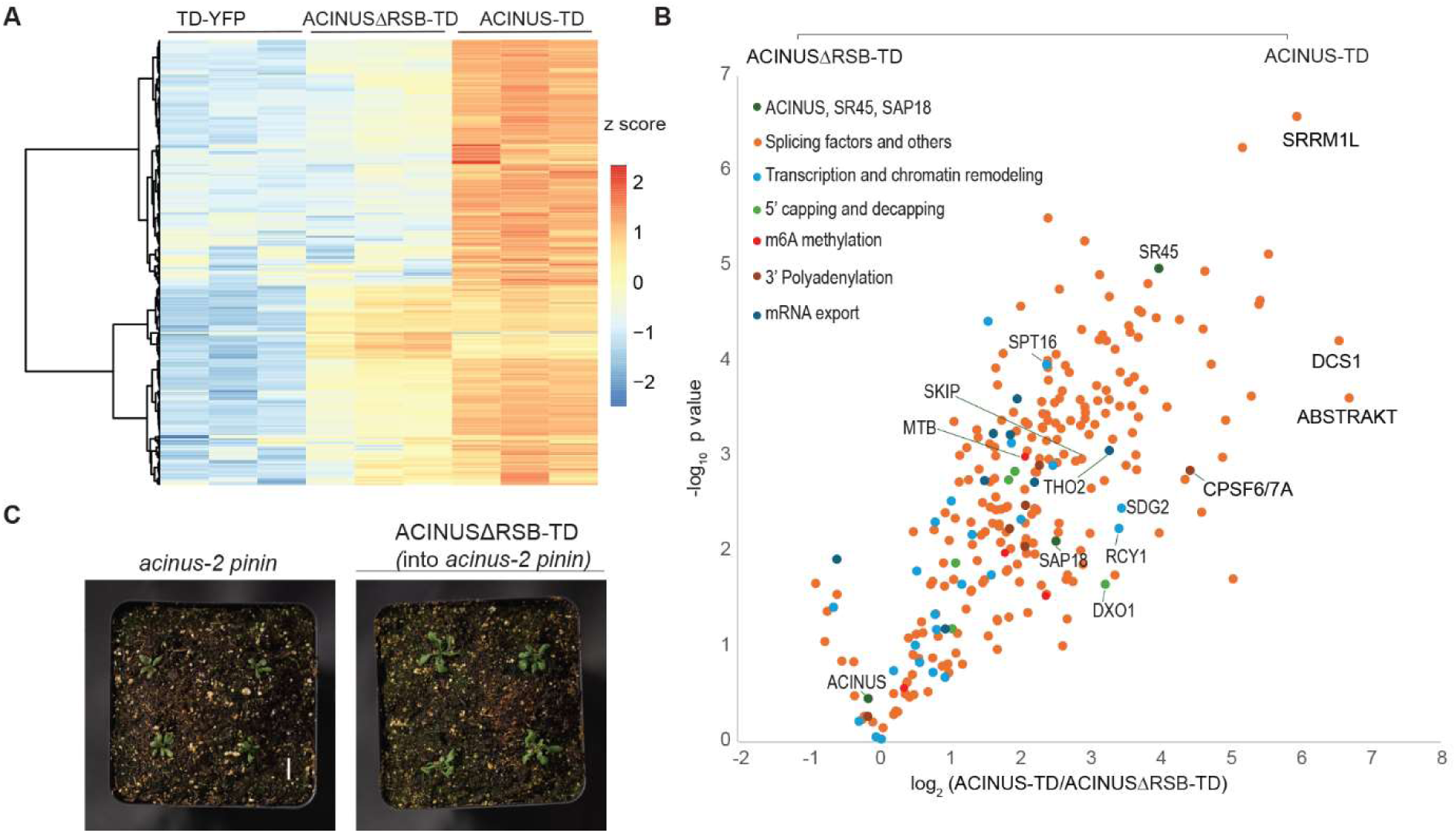
The RSB domain is critical for ACINUS function as a hub protein. (**A**) Heatmap comparison of TD-YFP, ACINUSΔRSB-TD, ACINUS-TD showing that approximately 75% of ACINUS-TD interactors have reduced or lost interactions in ACINUSΔRSB-TD. (**B**) Classification of interactor proteins affected by RSB deletion, showing perturbed interactions not only with SR45 and SAP18, but also with most splicing factors, transcription and chromatin remodeling associated proteins, and other factors involved in mRNA processing. (**C**) Phenotypic analysis of the ACINUS-ΔRSB-TD transgenic line in the *acinus pinin* background shows increased plant size compared to the double mutant, while retaining most of the characteristic features.

When ACINUS-ΔRSB-TD was introduced into the *acinus pnn* double mutant, the transgenic plants showed a significant increase in leaf length, resulting in a larger plant size compared to the double mutant (Fig. 5C, and Fig. S5B-C). However, they retained the characteristic narrow, twisted rosette leaf, and small rosettes (Fig.5C), and delayed flowering phenotype (data not shown) of the double mutant. This suggests that disruption of interactions between ACINUS and proteins in the ACINUS proxiome may be the main cause of the striking phenotype.

### Mutation in known and novel interactors lead to overlapping molecular or morphological outcomes

We hypothesized that depletion of the proteins identified with the same network would result in similar or opposite phenotypic changes - either at the molecular level, such as AS events, or in morphological traits - compared to the *acinus pinin* double mutant. To test this, we first analyzed AS events in several mutants with available RNA-seq data (*skip, mac3a mac3b, prp18a, prp4ka, luc7a luc7b luc7rl, xrn3,* and *sac3a*) (54,58–61). We then evaluated the morphological phenotype in selected interactor mutants (*tgh*, *dxo1*) (47,62).

Among the validated proteins, SKIP, MAC3A and MAC3B were previously identified by ACINUS IP-MS. Both SKIP and MAC3A/MAC3B are components of the Nineteen Complex (NTC), which is essential for proper spliceosome assembly and activation (58). The other interactors are novel. PRP18A functions in the second step of pre-mRNA splicing (59), while PRP4KA is a serine/threonine kinase associated with the spliceosome machinery (54). The LUC homologs are part of the U1 snRNP (61), and XRN3 encodes a 5’ to 3’ exoribonuclease (60). These interactors show extensive connectivity within the ACINUS proxiome network, as indicated by the edge numbers (Fig. 6A and Fig. S4). In contrast, SAC3A (54), originally identified from an AS mutant screen using a splicing reporter, is a putative mRNA export factor, although its precise role in mRNA export remains to be elucidated (63). Notably, SAC3A has minimal connectivity with the network.

**Figure 6.**
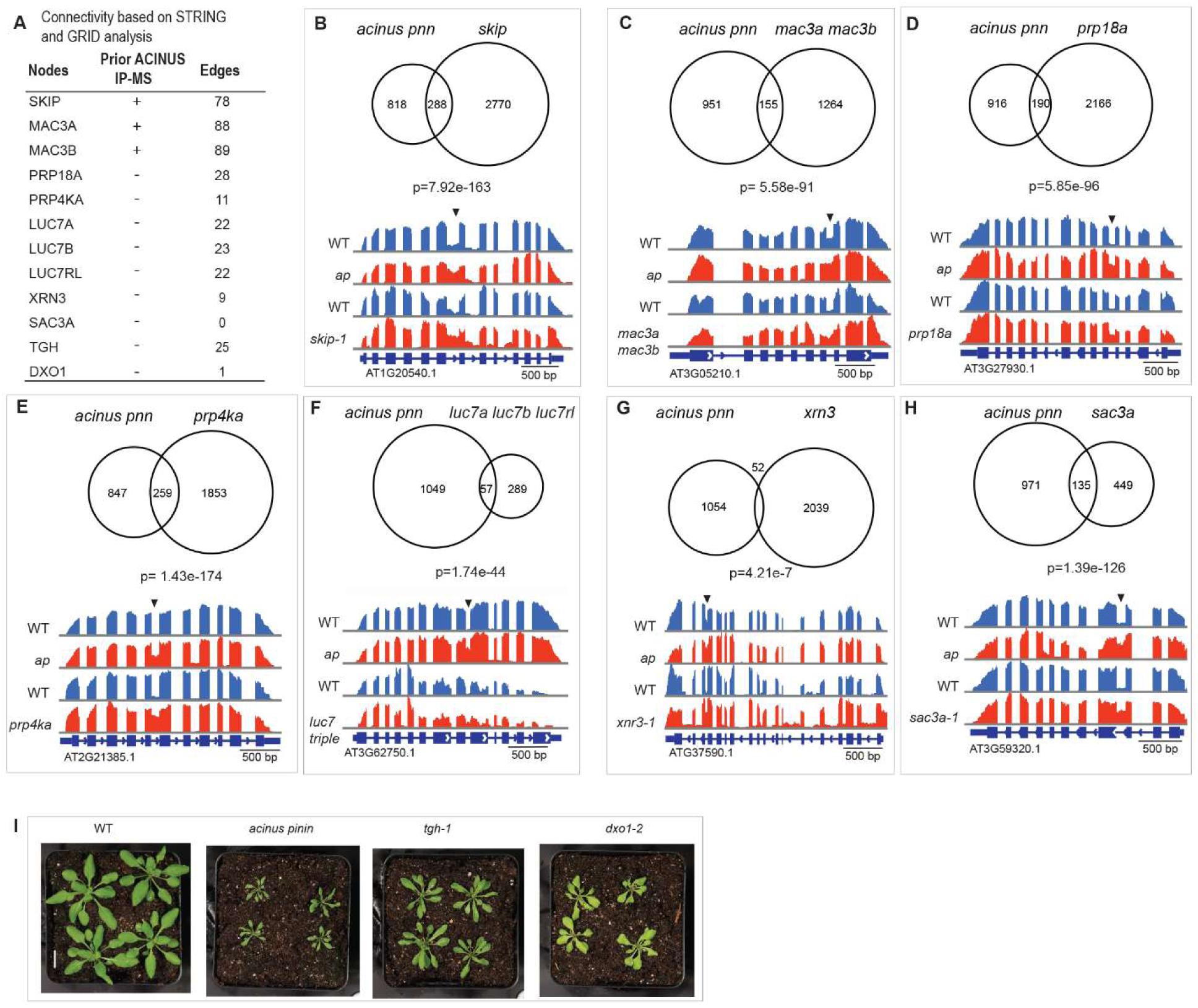
Mutations in known and novel interactors lead to overlapping molecular or morphological outcomes. (**A**) Network connectivity of selected interactors for validation, including three previously identified by ACINUS IP-MS and the rest as novel interactors. Some show extensive connectivity within the ACINUS proxiome network, as indicated by edge numbers, while others have minimal connections. (**B-H**) Venn diagram comparing common and distinct retained intron events between the *acinus pinin* mutant and several other mutants, with corresponding p-value. A representative retained intron event is shown for each comparison, with arrows indicating the retained introns. (**I**) *tgh* and *dxo1* mutants show phenotypic similarities to the *acinus pinin* double mutants, including twisted leaves, reduced size, and a pale green coloration.

We re-analyzed the RNA-seq results for the *skip, mac3a mac3b, prp18a, prp4ka, luc7a luc7b luc7rl, xrn3, and sac3a* mutants and compared the results with those of the *acinus pinin* mutants. Since retained intron is the dominant event in plant AS, we focused our analysis on this event. This analysis revealed a striking overlap in target events, with *skip* having the most shared events, and *xrn3* having the fewest (Fig. 6B-H, Fig. S6A). Surprisingly, ¼ of the retained intron events affected in the *sac3a* mutants were also affected in the *acinus pinin* mutants. Splicing factors can exert both positive and negative effects on AS in a context-dependent manner. In our analysis, most of the regulated events followed the same direction, suggesting that knockout of these components predominantly leads to increased intron retention (Fig. S6B).

We further evaluated two novel interactors, TGH and DXO1, by comparing the morphological phenotype. TOUGH (TGH) interacts with TATA-box binding protein 2 and colocalizes with the splicing regulator SR34 to subnuclear particles (62). DXO encodes a NAD-RNA decapping enzyme that activates RNMT1 to methylate the guanosine cap of mRNA (47,64), but its connection within the network is minimal. Comparing phenotypes, both *tgh* and *dxo1* mutants exhibited reduced plant size and narrow, often pointed and curled leaves, similar to but less severe than the phenotypes observed in the *acinus pinin* mutants. In addition, *dxo1* exhibited a pale green phenotype similar to that of *acinus pinin* mutants (Fig. 6I).

In particular, the involvement of XRN3 and DXO1 in 5’ capping and decapping, together with SAC3A as a putative mRNA transporter, indicates that factors involved in different steps of mRNA processing can collectively influence AS regulation. These findings confirm that both previously known and newly identified interactors contribute to the same molecular pathways, leading to overlapping molecular or morphological outcomes.

### SPY and SEC regulate transcription and alternative splicing through ACINUS-dependent and - independent mechanisms

Among the ACINUS network proteins, 47 were previously identified as O-GlcNAcylated, 30 were O-fucosylated, and 20 were doubly modified (Fig. 7A) (30–32), including ACINUS, HRLP, PRP16, SUA, CPL3 (Fig. 7B). We hypothesize that SPY and SEC may regulate gene expression via AS and/or transcription, both in an ACINUS-dependent and -independent manner. Previous qPCR analyses in *spy* and *sec* single mutants revealed splicing defects in several genes that are also affected in *acinus pinin* mutants (13). Therefore, we examined both single and double mutants to further understand these regulatory roles.

**Figure 7.**
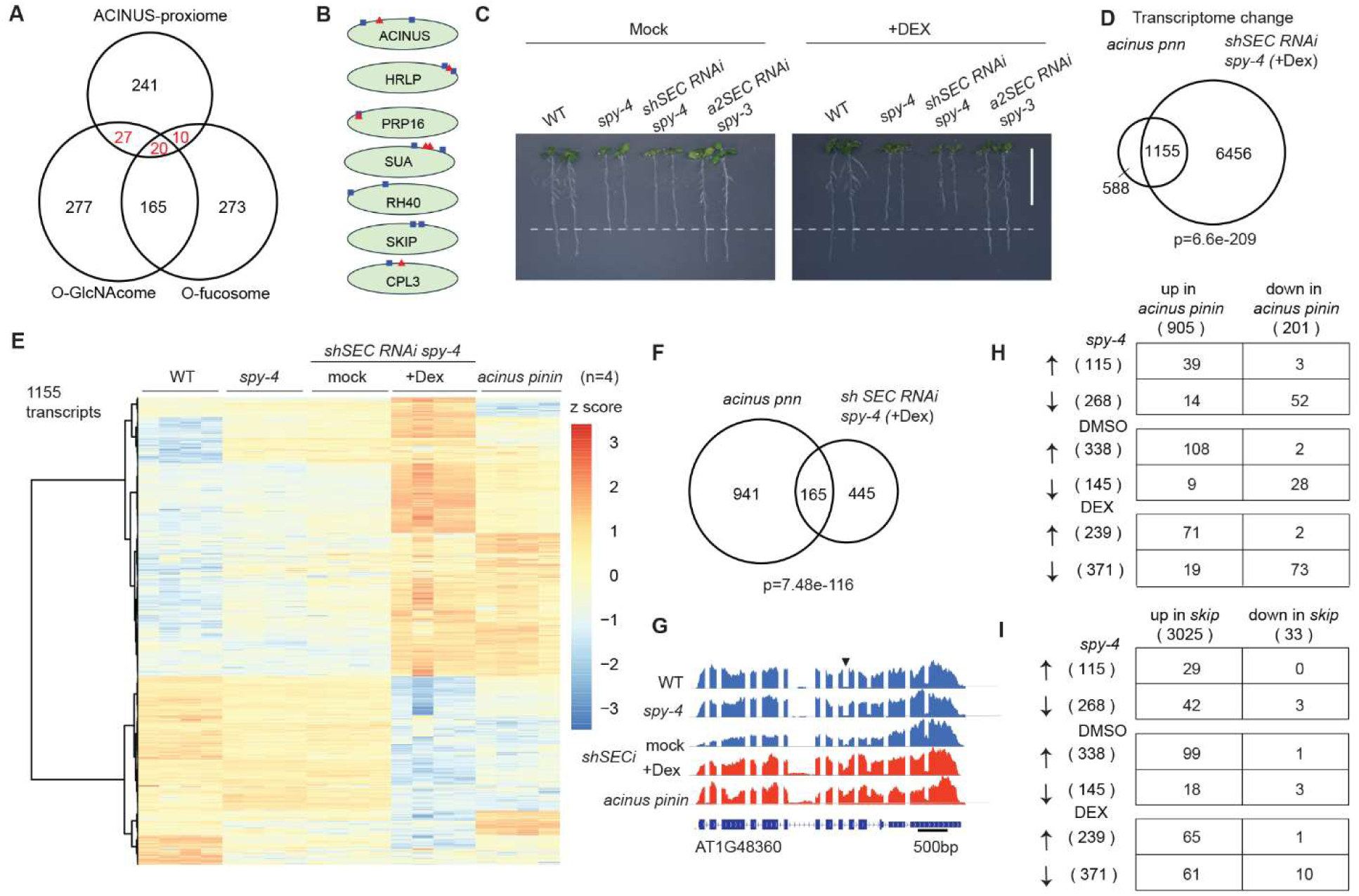
SPY and SEC regulate transcription and alternative splicing through ACINUS-dependent and -independent mechanisms. (A) Venn diagram showing the overlap of the ACINUS proxiome with the O-GlcNAcylation and O-fucosylation substrate libraries. (B) Selected proteins in the ACINUS proxiome that are O-GlcNAcylated (blue box), O-fucosylated (red triangle), or by both. (C) Inducible *shSEC RNAi* in *spy-4* showed reduced root growth after Dex treatment. (D) Significant overlap of transcriptome changes between *acinus pinin* and *shSEC RNAi spy-4* (+Dex) compared to WT. (E) Heatmap of 1,155 common transcripts between WT, *spy-4*, *shSEC RNAi spy-4* under mock or Dex treatment. (**F**-**G**) Significant overlap of retained intron changes between *acinus pinin* and *shSEC RNAi spy-4* (+Dex) compared to WT (F), with an example of increased retained 9th intron of AT1G48360 (G). (H) Regulated retained intron events occur predominantly in the same direction by ACINUS/PININ and SPY/SEC. (I) Comparison of retained intron changes between *skip* and *spy sec* double mutant reveals some SKIP-dependent manner.

To elucidate the role of SPY and SEC in AS masked by embryonic lethality (31,65), we generated a conditional double mutant by introducing short hairpin *SEC RNAi* into a *spy-4* null background (termed as *shSECi*) (Fig. S7A). This line resembled *spy-4* on control plates with short roots, but exhibited early seedling lethality on plates containing dexamethasone (Dex)(Fig. S7B). When *shSEC RNAi spy-4* seedlings were transferred from normal plates to Dex-containing plates, they showed progressively reduced root growth by day 3, leaf discoloration by day 5, and complete growth arrest by day 7 (Fig. 7C and Fig. S7C-D). These phenotypes are similar to, but more severe than, those observed in the *a2 SEC RNAi spy-3* inducible line (66) (Fig. S7C-D).

We performed a genome-wide comparison of AS and transcription in WT, *spy-4*, *acinus pnn*, and *shSECi* lines (see Methods). A *sec* single mutant was excluded due to the lack of phenotype, which is likely compensated by SPY (67). To assess data quality and to evaluate the potential effects of DMSO treatment, we performed a principal component analysis (PCA) on the transcriptomic data. The PCA results confirmed that replicates from WT, *acinus pinin,* and Dex treated *shSECi* clustered closely together, and that their biological groupings are distinct (Fig. S8A). The biological groupings of DMSO treated *shSECi* and *spy-4* lines clustered closely together, which indicates that DMSO has minimal transcriptional effects on seedlings. A total of 1,155 up- or down-regulated transcripts were shared between *acinus pnn* and Dex treated *shSECi* seedlings (log_2_FC ≥ 2 and FDR≤ 0.05), accounting for 66.3% of *acinus pnn*-dependent and 18.6% of *shSECi* (+Dex)-dependent events, when compared to WT (Fig. 7D-E, Fig. S8B).

Next, we analyzed the retained intron changes shared by the *acinus pnn* and *shSEC RNAi spy-4* (+Dex) mutants. Remarkably, 165 retain intron events were common to both mutants, accounting for 14.9% of *acinus pnn*-dependent events and 27.0% of *spy sec-*dependent events (Figure 7F-G). Co-dependent splicing events tended to be similarly altered in *acinus pnn* and *shSEC RNAi spy-4* (+Dex), with either increased or decreased intron retention (Fig. 7H). Comparable results were also observed in the *spy-4* mutant and *shSEC RNAi spy-4* (+ mock) (Fig. 7H), with the latter showing surprisingly more increased intron retention events. In contrast, a comparison with the *skip* mutant revealed a significant overlap of events, but mainly for increased intron retention (Fig. 7I); SKIP has previously been shown to be O-GlcNAc modified (Fig. 7B). These common and distinct IR and transcription events suggest that SPY and SEC regulate AS and transcription in both an ACINUS-dependent and -independent manner, highlighting the complexity of plant regulatory systems.

## Discussion

Significant efforts have been made to map the repertoire of proteins involved in AS using various proteomic and genetic approaches, including IP-MS with splicing factors or unspliced mRNA as bait, systematic RNAi or CRISPR-Cas-mediated gene ablation, and splicing reporter markers, as recently reviewed by Ule and Bencowe (20). However, these methods often lack the sensitivity to detect transient or weakly associated interactors and are prone to false negatives due to the high redundancy observed in animals. Our TurboID-based analysis overcomes these limitations and provides an exceptionally powerful method for identifying both known and novel components involved in AS. By integrating label-free quantification (LFQ) and SILIA-^15^N with TurboID-based LC-MS/MS, we enriched proteins at both the protein and peptide levels for ACINUS-TD, while also incorporating PININ-TD and SR45-TD, generating the first comprehensive map of the ACINUS-centered network regulating AS. Notably, we identified 35 interactors previously detected by ACINUS IP-MS, but more importantly, we identified over 260 novel interactors, demonstrating the unprecedented sensitivity of TurboID in the discovery of ACINUS associated proteins. Many of these interactors show reduced or abolished interaction with ACINUS upon deletion of the RSB domain. This high-resolution proximity labeling approach allows for a more reliable and comprehensive characterization of the plant AS machinery, capturing highly transient interactors such as kinases and methyltransferases known to regulate AS, as well as multiple members of the same protein families. The observed redundancy highlights the complexity of the AS regulatory systems in plants, mirroring the functional plasticity and complexity of the animal splicing machinery, and is consistent with the biochemical and cryo-electron microscopy studies showing that eukaryotic spliceosomes, such as the human spliceosome, contain a greater number of regulatory components and are significantly more dynamic than the yeast spliceosome (68,69).

Interdependent mRNA processing is well established in both mammals (70) and plants (71), with evidence that the C-terminal domain of RNA polymerase II (RNAPII) recruits splicing factors to nascent transcripts. Long-read sequencing of nascent RNA chains further supports this coupling, revealing a tightly coordinated integration of alternative splicing (AS) with transcription. Consequently, AS is dynamically influenced by transcriptional kinetics and chromatin architecture, and vice versa. Our ACINUS proxiome analysis identified numerous transcriptional and chromatin remodeling factors that support the co-transcriptional nature of AS. Furthermore, the detection of proteins involved in 5’ capping and decapping, m6A methylation, 3’ cleavage and polyadenylation, and mRNA export strongly suggests that these processes are intricately linked to AS regulation (Fig. 8A).

**Figure 8.**
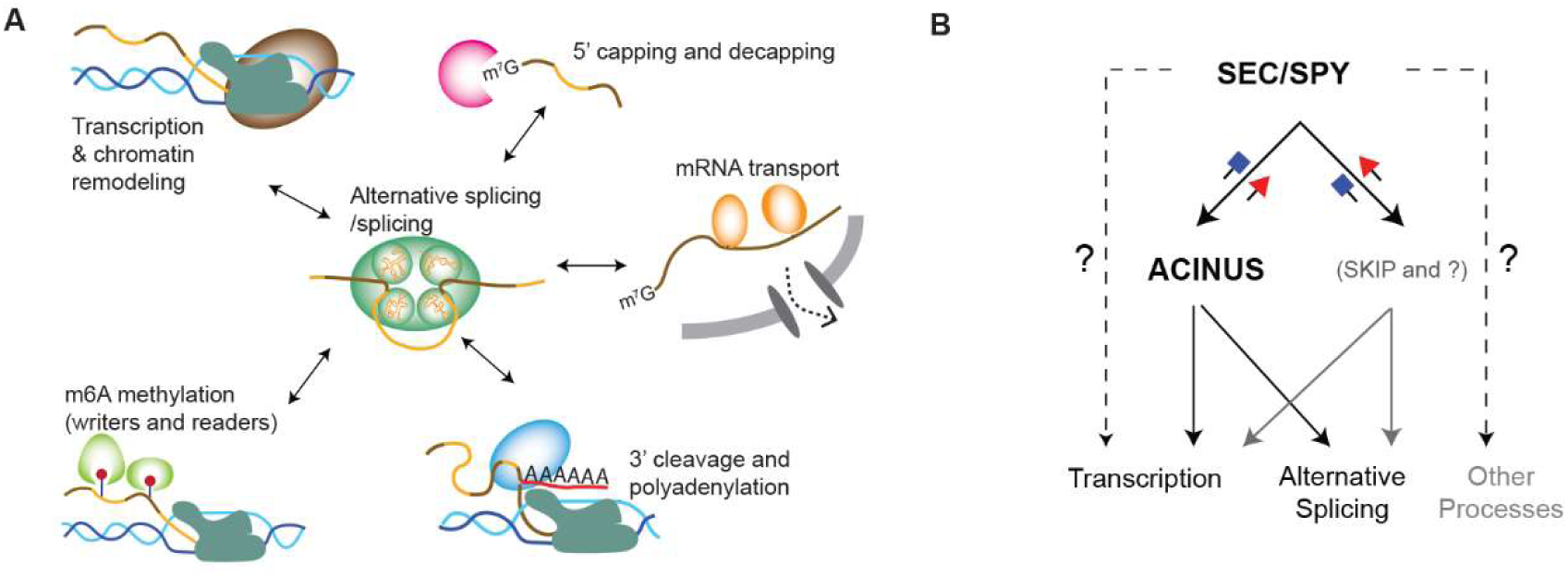
Model of interdependent mRNA processing and the role of SEC/SPY in regulating alternative splicing and transcription through ACINUS-dependent and -independent pathways. (A) Evidence from ACINUS-proxiome profiling suggests that alternative splicing occurs within large macromolecular complexes and is tightly coordinated with other mRNA processing events, including transcription, chromatin modeling, 5’ capping and decapping, m6A methylation, 3’ cleavage and polyadenylation, and mRNA transport. RNA-seq analysis further demonstrates that mutations in these mRNA processing pathways affect AS outcomes, highlighting the interdependence of these processes. The detection of enzymes such as kinases and methyltransferases in the ACINUS-proxiome suggests conserved mechanisms of AS regulation across species, while many ACINUS-associated factors exhibit plant-specific properties. These reveal both conserved and unique regulatory landscape of AS in plants. (B) RNA-seq data suggest that SEC and SPY modulate transcription and alternative splicing through both ACINUS-dependent and ACINUS-independent mechanisms.

Functional validation of interactors further supports this interconnected model. RNA-seq analysis and morphological characterization of interactor mutants provide compelling evidence that both previously known and newly identified ACINUS-associated proteins co-regulate common molecular pathways, leading to overlapping phenotypic outcomes. For example, a recent study on RNA polymerase II C-terminal domain phosphatase-like 3 (CPL3) shows a key regulatory role in AS, preferentially affecting the splicing patterns of defense genes(72). Our studies show that CPL3 is associated with the ACINUS proxiome.

Furthermore, the involvement of XRN3 and DXO1 in 5’ capping and decapping, together with SAC3A as a putative mRNA transporter, highlights a critical and previously underappreciated interplay between distinct mRNA processing steps in shaping AS outcomes. This underscores the extensive cross-talk between different mRNA regulatory mechanisms and their coordinated influence on alternative splicing.

The supraspliceosome hypothesis proposes that mRNA processing occurs within phase-separated condensates that spatially and temporally coordinate multiple processing steps to ensure efficient and regulated mRNA maturation(20,73–75). Our results provide strong evidence that such a mechanism exists in plants, demonstrating that AS is not an isolated event but is embedded in a highly interconnected network.

The conserved and species-specific features of AS, in particular the dominance of intron retention in plants versus exon skipping in animals (29,76), remain puzzling, in particular, especially with respects to how the AS machinery recognizes binding sites and assembles to define introns and exons. Our bioinformatic analysis, incorporating STRING and GRID data, BLAST for homolog searches, and IntroPro domain analysis, reveals both conserved and divergent mechanisms between plants and animals. The detection of numerous homologs in animals, along with shared domain features and the identification of kinases and methyltransferases, suggests conserved functions. In contrast, plant-specific features, such as 41 proteins without human homologs and the detection of SR proteins with plant-specific subfamilies, may explain why plants exhibit different forms of AS despite overall similarities in the composition of the AS machinery between plants and animals.

Furthermore, we show that SPY and SEC regulate AS and transcription through ACINUS-dependent and -independent pathways (Fig. 6B). Our RNA-seq data shows that SPY and SEC regulate over 7000 gene expressions and over 600 intron retention events. To distinguish between direct and indirect effects, we performed RNA-seq analysis 3 days after Dex treatment, a time point when the conditional double mutants have detectable RNAi expression and are just beginning to show phenotype. At this early stage, it is possible that SEC is not fully inhibited, leaving residual SEC activity that could influence the observed molecular phenotype. Endogenous SEC expression is hard to detect by mass spectrometry (77), making the quantitative analysis of SEC protein expression in the conditional double mutant difficult. Nevertheless, the clear pattern of intron retention provides strong evidence that SPY and SEC regulate alternative splicing and transcription through both ACINUS-dependent and -independent pathways. We also surprisingly observed more increased intron retention in DMSO-treated plants, while transcriptional analysis shows that they are similar to *spy-4* single mutants, it is possible that there is some leaky expression, and that SPY and SEC have opposite and overlapping roles (78,79).

Our study shows that sugar modifications have profound effects on AS and transcription. Future research should focus on understanding how sugar modifications specifically affect the splicing machinery to influence its functions. Additional genetic, biochemical, and structural studies, including a more comprehensive analysis of the AS components identified in our study, are needed to fully elucidate the role of these proteins in AS, uncover the mechanistic basis of their involvement in RNA processing, and explore the evolution of AS.

## Acknowledgement

We thank Dr. Laila Moubayidin lab at John Innes Center for sharing the a2 SECamiRNAi line, Dr. Xiao-Ning Zhang at St Bonaventure University for sharing *sr45-1* mutant seeds, and Dr. Andrea Mair and Dr. Dominique Bergmann at Stanford University for providing the TurboID gateway clones, and Daniel Wang for bioinformatics analysis. We also thank the Arabidopsis Biological Resource Center (ABRC) for providing several mutant seed stocks and to William Conner for photographing mutant phenotypes. This work is supported by the National Institutes of Health (R01GM135706 and S10OD030441to S.-L.X., and diversity supplement to A.V.R.; and R01GM066258 to Z.-Y,W), Carnegie Endowment Fund to the Carnegie Mass Spectrometry Facility.

## Contribution

R.S, D.B, A.V.R and S.K. generated TurboID constructs and transgenic lines. R.S. carried out TurboID experiments. A.V.R performed data mining and refined the IR analysis pipeline with assistance from W.D.L. S.K. generated and characterized *shSECi* lines in *spy-4* and *spy-3* line under supervision by S.L.X and Z.Y.W. S.C. and S.K. generated RNA-Seq data and S.C. and A.V.R analyzed the RNA-seq data. J.L. and H.C. generated shSECi construct. S.K. and K.A. performed microscopy, W.Z. characterized truncated construct phenotype. S.L.X. conceived the projects and wrote the manuscript.

## Data Availability

The mass spectrometry proteomics data have been deposited to the ProteomeXchange Consortium via the PRIDE partner repository with the dataset accession numbers: PXD059672 and PXD059673. The RNA-seq data have been deposited at NCBI Gene Expression Omnibus (GEO) database under the access number GSE286870. All other data is available from the corresponding author on reasonable request.

## Methods and materials

### Plant materials and growth conditions

All Arabidopsis plants used in this study were in the Columbia-0 (Col-0) background. The *tgh-1* mutant (SALK_053445) and the *dxo1* mutant (SALK_103157) were obtained from the Arabidopsis Biological Resource Center (ABRC). For seedlings grown on the medium in petri dishes, all the sterilized seeds were stratified at 4°C for 2-3 days and grown on the specified medium for the time indicated in each experiment.

For plants grown in the greenhouse, a 16-h light/8-h dark cycle at 22–24 °C was used for general growth or photo collection, as described in each experiment. Most of the seedlings were grown vertically on ½ Murashige and Skoog medium and supplemented with 0.8% (w/v) phytoagar under constant light (24 h light, 22°C) for the time indicated in each experiment. For protein-level enrichment of biotinylated proteins using label-free quantification (LFQ), seedlings were harvested after 10 d and treated with biotin at 50 µM for 1 h. For peptide-level enrichment of biotinylated proteins using SILIA, the bait and control lines were grown at 22°C on ^14^N or ^15^N medium plates (Hogland’s No2 salt mixture without nitrogen 1.34g/L, 6g/L phytoblend, and 1g/L KNO_3_ or 1g/L K^15^NO_3_ (Cambridge Isotope Laboratories) for ^14^N medium or ^15^N medium, respectively, pH5.8) for 14 days and then treated with 50 µM biotin for 1h.

For RNA-seq, wild-type (WT), *acinus pinin*, *spy-4* were grown on ½ MS for 8 days before harvesting. *shSEC RNAi spy-4* were grown on ½ MS plates for 5 days and transferred to mock (DMSO) or Dex (10µM)-supplemented plates for 3 days before harvesting. For phenotypic analysis, the same conditions were used, but phenotypes were further observed after 5 and 7 days of transfer. Similar lines were also used for phenotypic analysis directly on mock or Dex-treated plates, where the plants were grown on ½ MS medium for 10 days on either mock (DMSO) or Dex (10µM in DMSO) plates. All plants were grown under constant light.

### Construct generation and transformation

The full-length cDNA of ACINUS, PININ and SR45, and the truncated cDNA of ACINUSΔRSB were cloned into pENTRD-TOPO vectors. The promoters, approximately 2 kb upstream of the ATG start codon of ACINUS, PININ and SR45, were amplified and cloned into the pENTR5 vector gateway clone to generate the promoter entry clone (pENTR5-ACINUSpro, or PININpro, or SR45pro). The ACINUS-TD, PNN-TD, SR45-TD, and ACINUSΔRSB-TD constructs were generated by multisite gateway cloning, allowing LR reactions of three entry vectors (the promoter entry clone, the cDNA entry clone, and pDONR_P2R-P3_R2-Turbo-mVenus -STOP-L3) with the destination vector pB7m34GW (41) to generate the final expression vector. The control constructs TD-YFP under three different promoters were generated by multisite gateway cloning, allowing LR reactions of two entry vectors (the promoter entry clone, pENTR_L1-Turbo-YFP-NLS-STOP-L2) with the destination vector R4pGWB501 to generate the final expression vector. Related primers are listed in Supplementary Table S2.

ACINUS-TD, PININ-TD were transformed into the *acinus-2 pinin* mutants, SR45-TD was transformed into the *sr45-1* mutants. ACINUSΔRSB was transformed into WT for TurboID experiments, and into *acinus-2 pinin* mutants for phenotypic analysis. These lines were selected using BASTA. The various TD-YFP controls were transformed into wild-type and selected on hygromycin. Complemented bait-TD lines and TD-YFP lines were selected for their expression level or complementation phenotype or both. T3 lines were used for phenotypic analysis and TurboID experiments.

*shSEC RNAi* lines: The short hairpin SEC RNA (*shSEC RNAi*) constructs for *SEC* were made by linking two fragments PCR-amplified from the 5’ end of the genes (genomic DNA or gDNA, 34-846 relative to ATG start site) and cDNA (34-389 relative to ATG start site) in an head-to-head orientation, with primers specified in primer table (SEC_gDNAF/SEC_R to amplify genomic DNA; SEC_cDNA_F/SEC-R to amplify cDNA). XhoI or PstI site was added to the 5’ end of the primers used for cloning. The inverted constructs were first cloned in pBluescript II SK+ and then subcloned into the XhoI and SpeI sites in the Dex-inducible binary vector pTA7002(80). The construct was transformed into the *spy-3* and *spy-4* mutant. Over 100 T2 independent lines in the *spy-3* were screened, with 2 lines showing reduced root phenotype. In the *spy-4* background upon Dex treatment, a total of 14 independent lines were screened, and 4 lines showed a strong growth arrest phenotype in the presence of Dex treatment. These T3 homozygous lines were used for phenotypic analysis and RNA-seq.

### Functional analysis of bait-TD fusion complementation analysis

The following lines were sterilized and grown on ½ MS plates under constant light for 10 days: WT, *acinus-2 pinin*, *sr45-1* mutant, ACINUS-TD/*ap*, PININ-TbID/*ap*, SR45-TbID/*sr45-1*, then transplanted to soil and grown under long day conditions (L/D of 16h/8h) at 22°C. Photos were taken of plants at approximately 3 weeks old and 5 weeks old to compare leaf size and shape, and flowering time. Similarly, *acinus-2 pinin,* ACINUSΔRSB-TD/*ap* were grown on ½ MS plates under constant light for 7 days then transplanted to soil and grown under long day conditions (L/D of 16h/8h) at 22°C. Photos were taken at plants of approximately 3 weeks old to compare leaf size and shape.

### Confocal Microscopy and Spinning disk Microscopy

Images of 4-day-old Arabidopsis seedlings expressing BAIT-TurboID (which also contains VENUS) constructs were taken with a Leica SP8 confocal microscope and spinning disk microscope. Images were processed using Adobe photoshop and ImageJ software.

### Root length quantification assay

Wild-type (WT), *spy-4*, *shSEC RNAi spy-4*, and *a2SEC RNAi spy-3* seeds were grown on ½ MS medium supplemented with 10 μM Dex or DMSO. After 10 days, the seedlings were photographed for phenotypic analysis to assess their response to Dex treatment. Alternatively, seeds were grown on ½ MS plates for 5 days before being transferred to ½ MS medium containing either 10 μM Dex or DMSO mock treatment. Seedlings were imaged at 0-, 3-, 5-, and 7-days post-transfer, and root elongation was measured using ImageJ software. Over 15 seedlings per sample were measured.

### Biotinylated protein enrichment at protein-level and peptide-level

Tissue samples (1 g each) were harvested in triplicate per condition. Biotinylated proteins were enriched at the protein-level using MyOne Streptavidin C1 Dynabeads (Invitrogen) with steps described in (41). The ACINUS-TurboID experiments included WT, ACINUS-TD, ACINUSΔRSB-TD/WT, and TD-YFP. The PNN-TurboID experiments included PNN-TD and a corresponding TD-YFP control, while the SR45-TurboID included SR45-TD and its TD-YFP control.

For peptide-level enrichment, ∼ 2 g of tissue was harvested per sample, with 4 biological replicates (two ^14^N-labeled and two ^15^N labeled) for both bait and control, Tissues were equally mixed as the following: Fw1: ^14^N ACINUS-TD /^15^N YFP-TD; Rv1, ^15^N ACINUS-TD/ ^14^N YFP-TD; Fw2: ^14^N ACINUS-TD /^15^N YFP-TD; Rv2:^15^N ACINUS-TD/ ^14^N YFP-TD. Proteins were extracted from mixed tissues using phenol extraction, digested and cleaned as described as (30). The peptide mixture was resuspended in the PBS buffer and incubated with 100 μL of Dynabeads M-280 Streptavidin (Thermo, 11206D) prewashed in the PBS buffer. The resulting mixture was incubated for 1 h at room temperature with shaking at 800 rpm. The beads were washed first with PBS twice, then with PBS with 5% ACN twice and finally with ddH_2_O once, each for 5 min at room temperature and 800 rpm. The biotinylated peptides were eluted with 200 μL of elution buffer (0.2% TFA, 0.1% FA, and 80% ACN) three times by heating at 75°C for 5 min. The eluted peptides were collected and dried by speedvac. The peptide mixtures were desalted using C18 OMIX Tips.

### LC-MS/MS analysis

Peptides were analyzed by liquid chromatography–tandem mass spectrometry (LC-MS) on an EasyLC1200 system (Thermo) connected to a high-performance quadrupole Orbitrap Q Exactive (Thermo), with parameters as described(41). Biotinylated peptides were eluted with a slight modification: from 3 to 6% solvent B (80% (v/v) acetonitrile/0.1% (v/v) formic acid) over 1 min, 6-10% solvent B over 4 min, 10 to 45% solvent B over 96 min, from 45 to 75% solvent B over 20 min, followed by a short wash of 8 min by 90% solvent B.

### Label-free Quantification for Protein-level enrichment and statistics processing using Perseus

Raw MS/MS data was loaded in Maxquant (v2.4.12). The search was using most of the preconfigured settings allowing match between runs options. Data was searched with a 4.5 ppm tolerance for precursor ions and 20 ppm for fragment ions. Peptide and protein FDRs were set as 0.01. Label-free quantification (LFQ) was enabled. Data was searched against the TAIR10 database *Arabidopsis thaliana* (https://www.arabidopsis.org/), concatenated with decoy protein sequences (a total of 35386 entries). A Maxquant LFQ was run with min ratio count as 2 (a minimum of 2 peptides quantification per protein) was required and match time window with 0.7 min, Quantification was done on unique and razor peptides and other parameters as default. Match between runs was applied. The data were filtered as described(41). A two-sided T-test with S0=2 and FDR=0.05 was used to determine significance and enrichment of protein.

### SILIA quantification for peptide-level enrichment using Protein Prospector

For Protein Prospector (version 6.4.9), MS/MS data were converted into peaklists using the script PAVA (peaklist generator that provides centroid MS2 peaklist). The peaklists were further filtered using signature ion peak 310.16 or 311.16 to generate filtered peaklists. Both full and filtered peaklists were searched using the same parameters, the former to confirm equal mixture of ^14^N- and ^15^N-labeled peptides and the latter to identify and quantify biotinylated peptides. A precursor mass tolerance of 10 ppm and MS/MS2 tolerance of 20 ppm were allowed. Carbamidomethyl (cysteine (C)) was searched for as a constant modification. Variable modifications allowed included biotinylation at uncleaved lysines (Biotin(K)), protein N-terminal acetylation, methionine (M) oxidation, N-terminal acetylation with M oxidation, N-terminal M-loss, N-terminal M-loss with acetylation, and peptide N-terminal Gln conversion to pyroglutamate, with three maximum variable modifications per peptide. Similar ^15^N parameters were used for ^15^N samples. Cleavage specificity was set to trypsin, allowing two missed cleavages. For Protein Prospector, default 1% FDR was allowed for peptides and 5% for proteins. The quantification was done as described in Shrestha et al. (42), (31) but reporting on both protein-level (medium ratio of quantified peptides from the same protein) and peptide-level (ratio of each peptide from the same protein). As ^14^N and ^15^N labeled peptides have different masses in both MS1 and MS2, peptides identified by both are of high confidence.

### Data integration and filtering

The data were rigorously filtered to generate a robust ACINUS-proxiome list (see Supplementary Table S1). The 1st category included 171 proteins (including three baits) that were enriched for all three baits: PNN-TD (protein-level) and SR45-TD (protein-level), and ACINUS-TD (protein-level or peptide-level). The 2nd included 42 proteins enriched by PNN-TD (protein-level) and ACINUS-TD (protein-level or peptide level). The 3rd category included 49 proteins enriched by SR45-TD (protein-level) and ACINUS-TD (protein-level or peptide-level). The 4th category identified 15 proteins that were enriched by ACINUS-TD (both protein-level and peptide level). The 5th included 21 proteins that are exclusively enriched by ACINUS-TD at the peptide level (in at least 3 reciprocally labeled experiments, ≥10 fold cut off).

### RNA Seq analysis and data mining

Paired-end RNA-seq data for WT, *spy-4*, *shSEC RNAi spy-4* (mock and Dex treated) samples were generated using the Illumina platform NovaSeq X Plus Series (PE150) at Novogene (California). mRNA libraries were prepared using polyA enrichment. Raw reads were trimmed using CutAdapt (Version 2.6), quality checked using fastqc (Version X) and aligned to the genome using STAR (Version 2.7.10b). STAR alignment was done with araport11 annotation (Araport11_GFF3_genes_transposons.Jan2024 and Araport11_GTF_genes_transposons.Jan2024) and TAIR10 genome sequence downloaded from arabidopsis.org. Alignment files were processed using RACKJ (Version 0.99a) to retrieve both read counts and read depth for each annotated intron and exon. For intron retention analysis the intron read depth ratio (IDratio) was calculated by dividing the intron read depth of the given intron by the average read depth of the two flanking exons (Fig. S1). However, for the first and last intron the IDratio is calculated using the depth of the second (3’) or first (5’) flanking exon, respectively, because read depths for the first and the last exon in a gene may be inaccurate due to sequencing bias. Undefined IDratios (e.g., when divided by zero) were imputed using a normal distribution downshifted by 1.8 standard deviations and a mean reduction by a factor of 0.3. All IDratios were log_2_ transformed prior to significance calculation (α < 0.05) in order to determine which introns were significantly increased in the *acinus pnn* double mutant. A two-tailed t-test was performed on both WT and mutant replicates (n =4). Introns were filtered using the following criteria: (1) p-value less than or equal to 0.05; (2) fold change greater than or equal to 1.5; (3) belonging to a gene with an average RPKM greater than 1; (4) average intron depth in the mutant is greater than or equal to 5; (5) not belonging to single intron genes; (6) mean IDratio of the mutant is greater than 0.15. To determine decreased retained introns, the average intron depth and the mean IDratio of the wild type was considered instead.

The calculation for exon skipping, alternative donor, alternative acceptor, alternative donor and acceptor were same as described in (13) with modified parameter by removing splicing events with average gene RPKM <1. P-value was set less than or equal to 0.05. The fold change cutoff was set as 1.5-fold.

For expression analysis, ReadsPerGene files from the STAR alignment were processed in R using Deseq2. The lfcShrink function was applied as described (81). Genes were defined as significantly differentially expressed if they had an adjusted p-value less than or equal to 0.05 and a fold change of less than or equal to 2. For all heatmaps, normalized gene counts were log_2_ transformed. All non-numerical values (e.g., -Inf) were imputed using the method described above. For PCA analysis, genes containing - Inf Values were removed.

For data mining, relevant RNA-seq data files were downloaded using NCBI sra-tool’s fasterq-dump (2.11.2) (Supplementary Table S1). The downloaded RNA-seq data was checked for quality using fastqc and subsequently processed by STAR aligner using the method described above. STAR parameters were adjusted for unpaired and/or shorter read lengths accordingly. Statistical significance for Venn diagram overlap was generated using a hypergeometric test in R. The analysis results were detailed in Supplementary Table 3.

## Supplementary Figures

**Supplementary Fig. S1.**
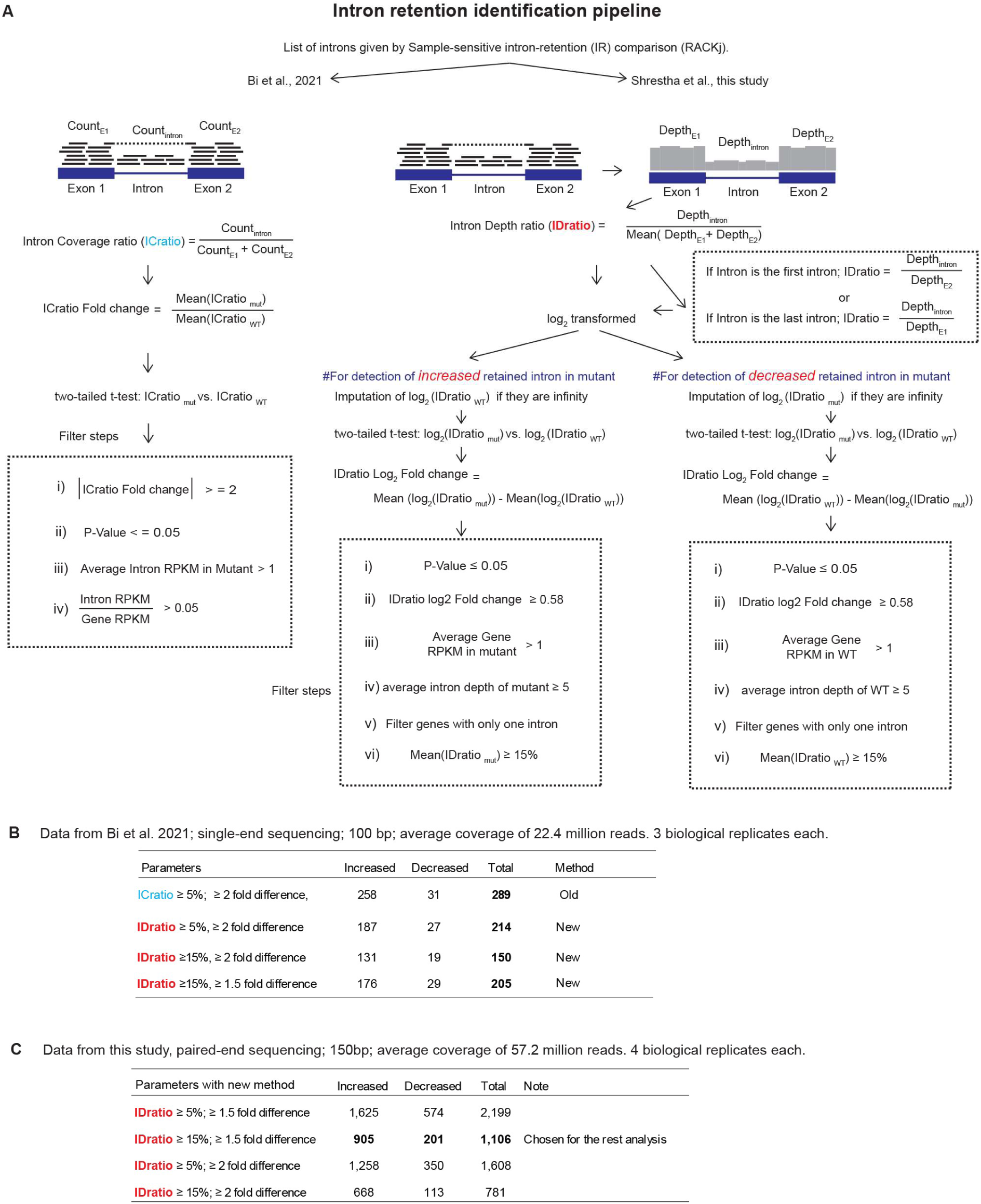
Improved pipeline for retained intron analysis. (A) Schematic of the improved intron retention analysis pipeline. The left panel illustrates the intron coverage ratio (ICratio) used by Bi et al., 2021, while the right panel introduces the intron depth ratio (IDratio). Key filtering steps are listed, including the application of more stringent thresholds, such as a mean IDratio of at least 15% in at least one group (WT or *acinus pinin*) for statistical analysis. (B) Comparison of parameters used for retained intron analysis with single-end sequencing data from Bi et al., 2021. The updated analysis pipeline applies stricter thresholds, reducing the number of detected altered retained intron events in *acinus pinin* double mutants. (C) Comparison of the detection of retained intron alternations in *acinus pinin* double mutants using new sequencing data and an updated analysis pipeline with different cutoff parameters. A higher IDratio was selected for further analysis, as a lower IDratio may lack biological significance despite statistical significance between WT and the double mutant.

**Supplementary Fig. S2:**
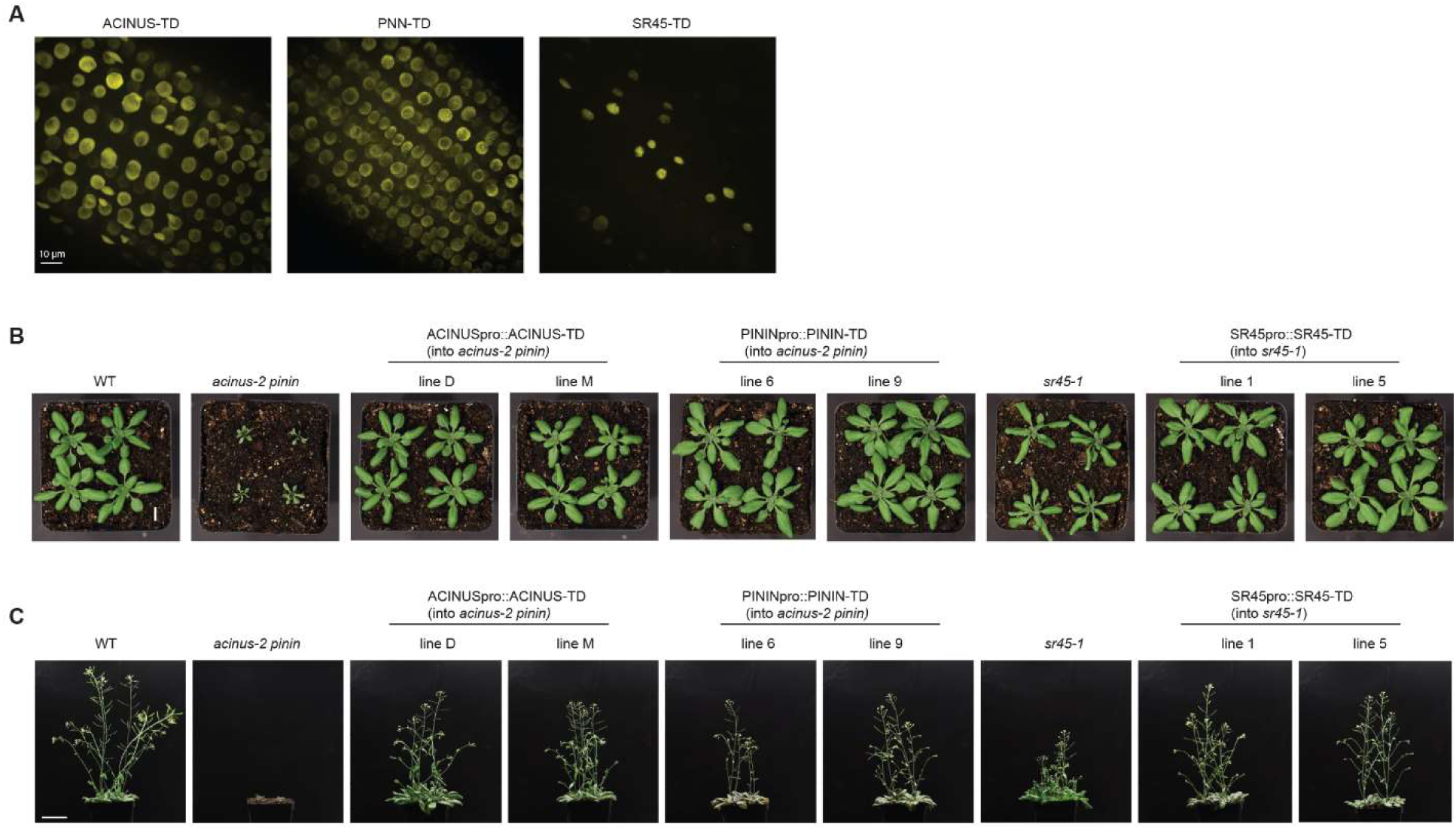
ACINUS-TD, PNN-TD, and SR45-TD fusion proteins localize to the nucleus and complement the leaf shape, rosette size, and flowering phenotypes in corresponding mutants. (A). Spinning disk images of ACINUS-TD, PNN-TD and SR45-TD. Scale =10 µm. (B). Wild type (WT), mutants, and two independent complementation lines of each bait were grown in soil for approximately 4 weeks, showing the complementation of the leaf shape and rosette size phenotypes. Scale = 1cm. (C). The same plants were grown in the greenhouse for about 6 weeks, showing the complementation of the flowering time phenotype by the TurboID fusion proteins. Scale bar=5 cm.

**Supplementary Fig. S3.**
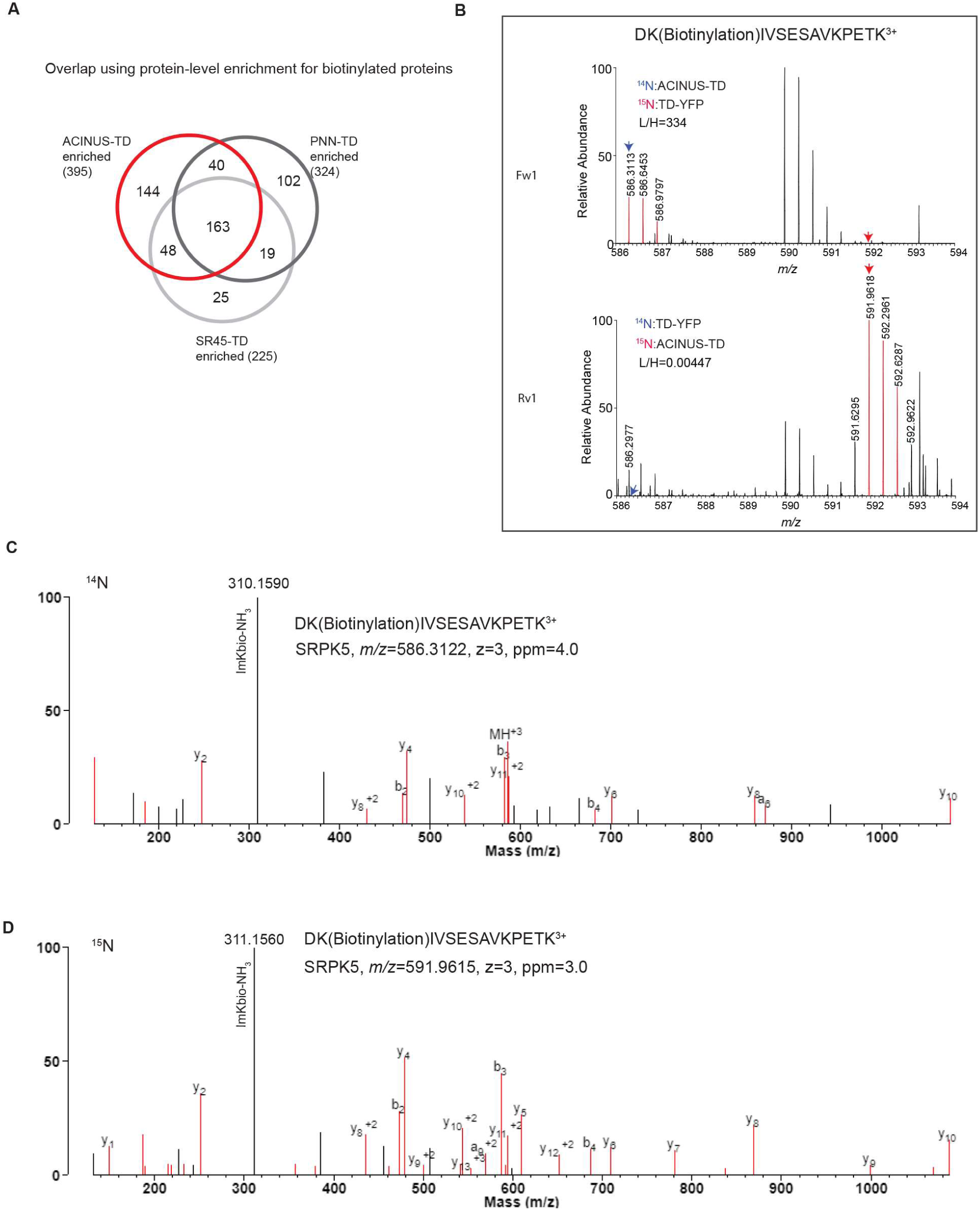
Overlap summary of enriched proteins and representative spectra for identification and quantification of SRPK5 biotinylated peptides. (A) Overlap summary of protein-level enrichment among three baits. (B) Quantification of SRPK5 biotinylated peptides from reciprocally labeled samples, with MS1 peak matches highlighted. Three peaks corresponding to M, M+1, M+2 peaks are highlighted in red for the upper ^14^N-labeled peaks and the lower ^15^N-labeled peaks from forward 1(Fw1) and reverse 1 (Rv1) experiments. Monoisotopic peaks are indicated by arrows, with ^14^N shown in blue and ^15^N in red. (**C**-**D**). MS2 spectra of *m/z* 586.3122 and 591.9615, 3+ precursors, identified a ^14^N-labeled (C) or ^15^N-labeled (D) biotinylated peptide from SRPK5 spanning from amino acid 208 - 221 with a modification at K209, detected in ACINUS-TD enriched samples. The signature ion corresponding to the derivative ion at *m/z* 310.16 (^14^N) or 311.16 (^15^N) due to the ammonia loss of the immonium ion of the biotinylated lysine (ImKBio, molecular formula C15 H19 N2 O2 S for ^14^N-labeled, and C15 H19 N 15N O2 S for ^15^N-labeled), is labeled with the measured mass.

**Supplementary Fig. S4.**
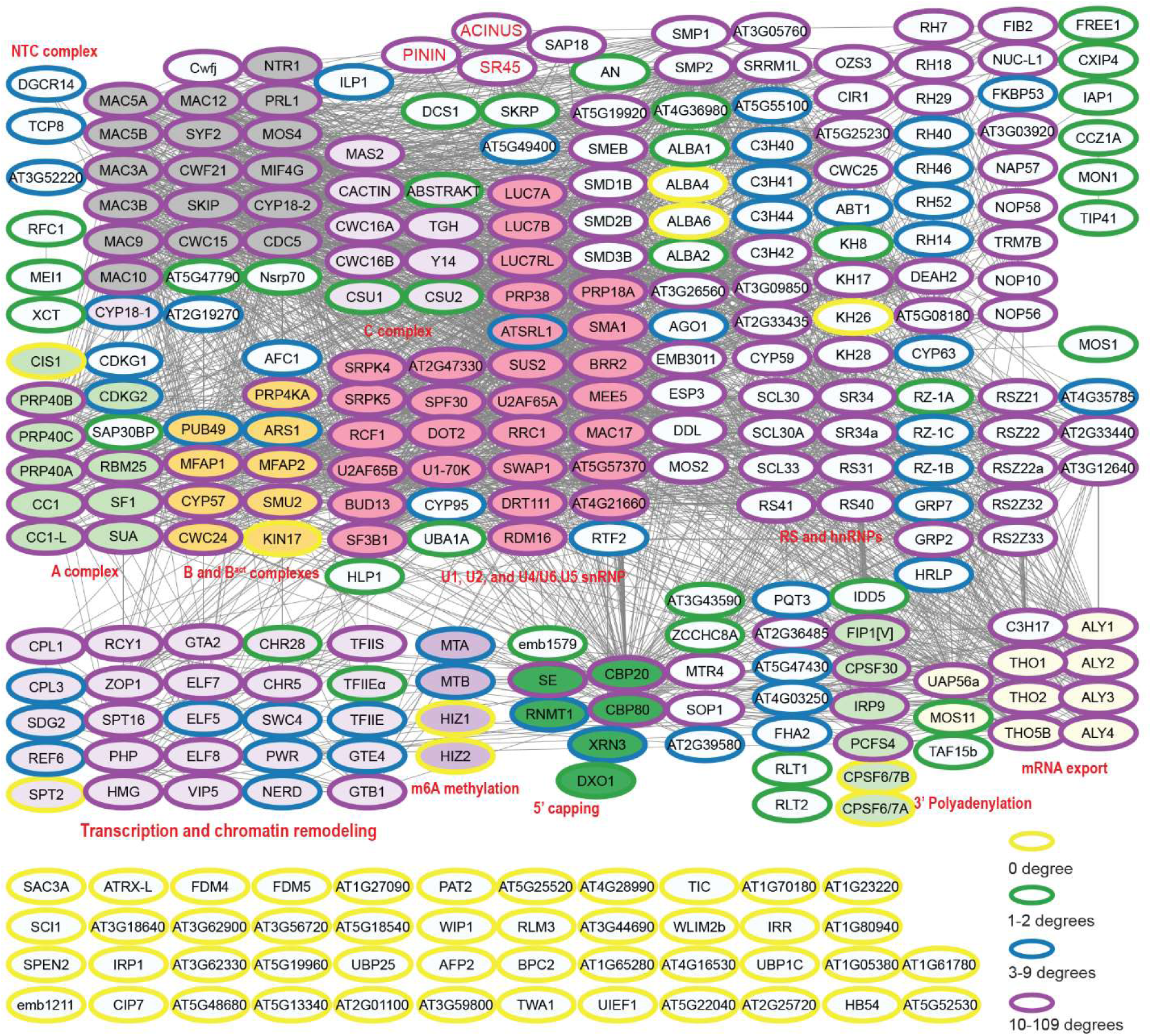
Cytoscape of the ACINUS-proxiome showing co-transcriptional, co-occurring activity of mRNA processing and high redundancy. The ACINUS-proxiome network is visualized in Cytoscape, integrating TurboID, STRING and GRID data. Nodes represent proteins and edges indicate interactions between them, as shown by gray lines. Proteins from STRING and GRID are circled based on connectivity: nodes with 10-109 edges are bordered in magenta, those with 3-9 degrees are bordered in blue, those with 1-2 edges are bordered in green, and those with no edges are bordered in yellow. Proteins with known or predicted functions are grouped and color-coded as follows: NTC complex filled in gray; A, B/B^act^, and C complex proteins are filled in green, yellow, and pink, respectively; core and accessory spliceosome components (U1, U2, U4/U5/U6) are filled in darker pink. Proteins involved in transcription and chromatin modeling are filled in light pink, while proteins involved in m6A methylation, 5’ capping, 3’ polyadenylation and mRNA export are filled in pink, dark green, light green, and light yellow, respectively.

**Supplemental Fig. S5.**
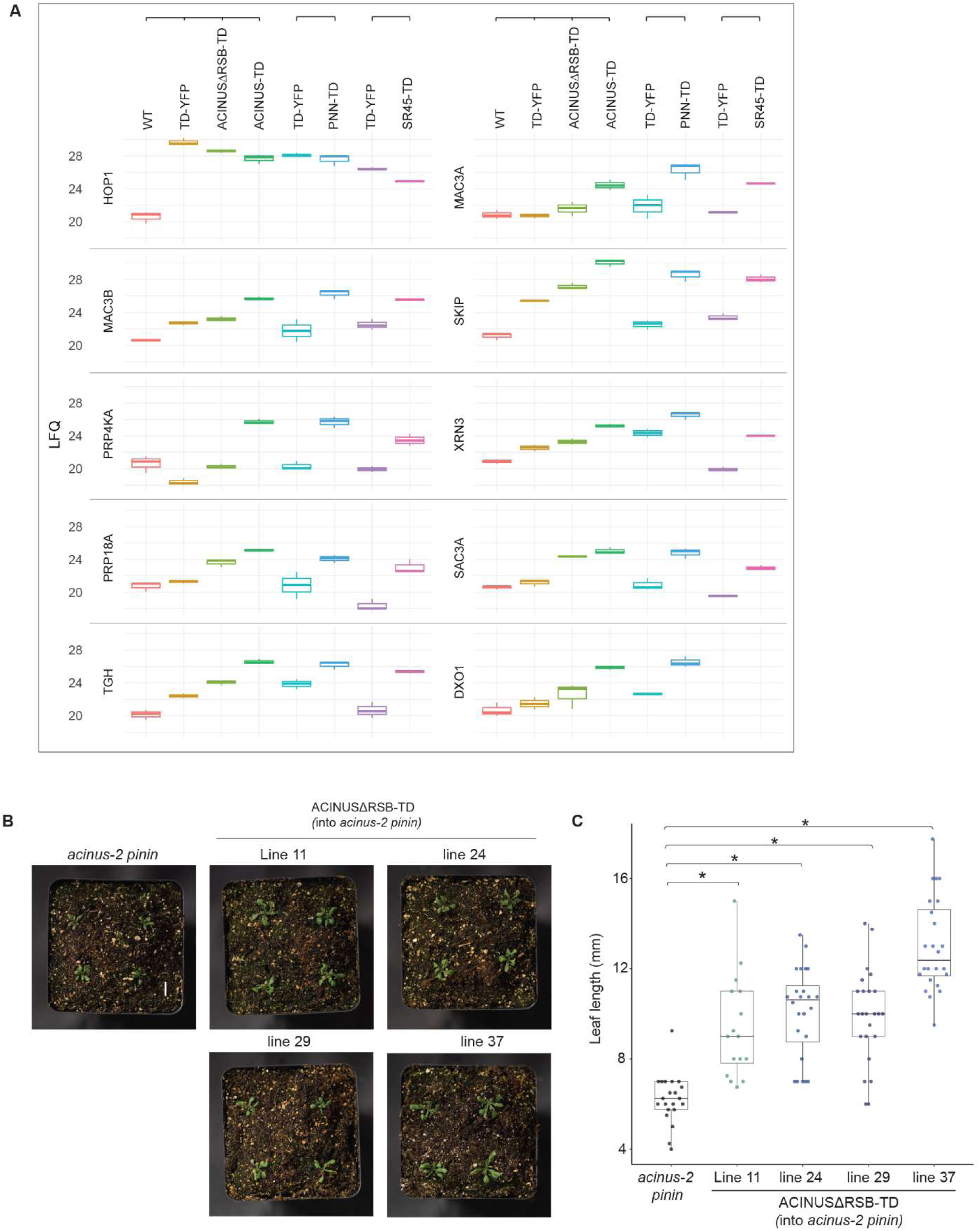
RSB deletion reduces or abolishes interactions and impairs ACINUSΔRSB-TD function in planta. (A) Quantification of selected proteins over multiple experiments shows reduced or lost interactions in ACINUSΔRSB. Selected examples include several ACINUS interactors and the nuclear standby protein HOP1, which is biotinylated by both baits and controls. The y axis represents the log2-transformed, normalized intensity of each protein across experiments. (B) Transgenic ACINUSΔRSB-TD lines in the *acinus pnn* mutant still show narrow and twisted leaves and overall small rosettes characteristic of double mutants. Four independent lines showing a consistent phenotype. (C) Quantification of leaf length of double mutant and transgenic plants shows that ACINUSΔRSB-TD expression results in larger leaves compared to the *acinus pnn* mutant. y axis shows leaf length.

**Supplemental Fig. S6:**
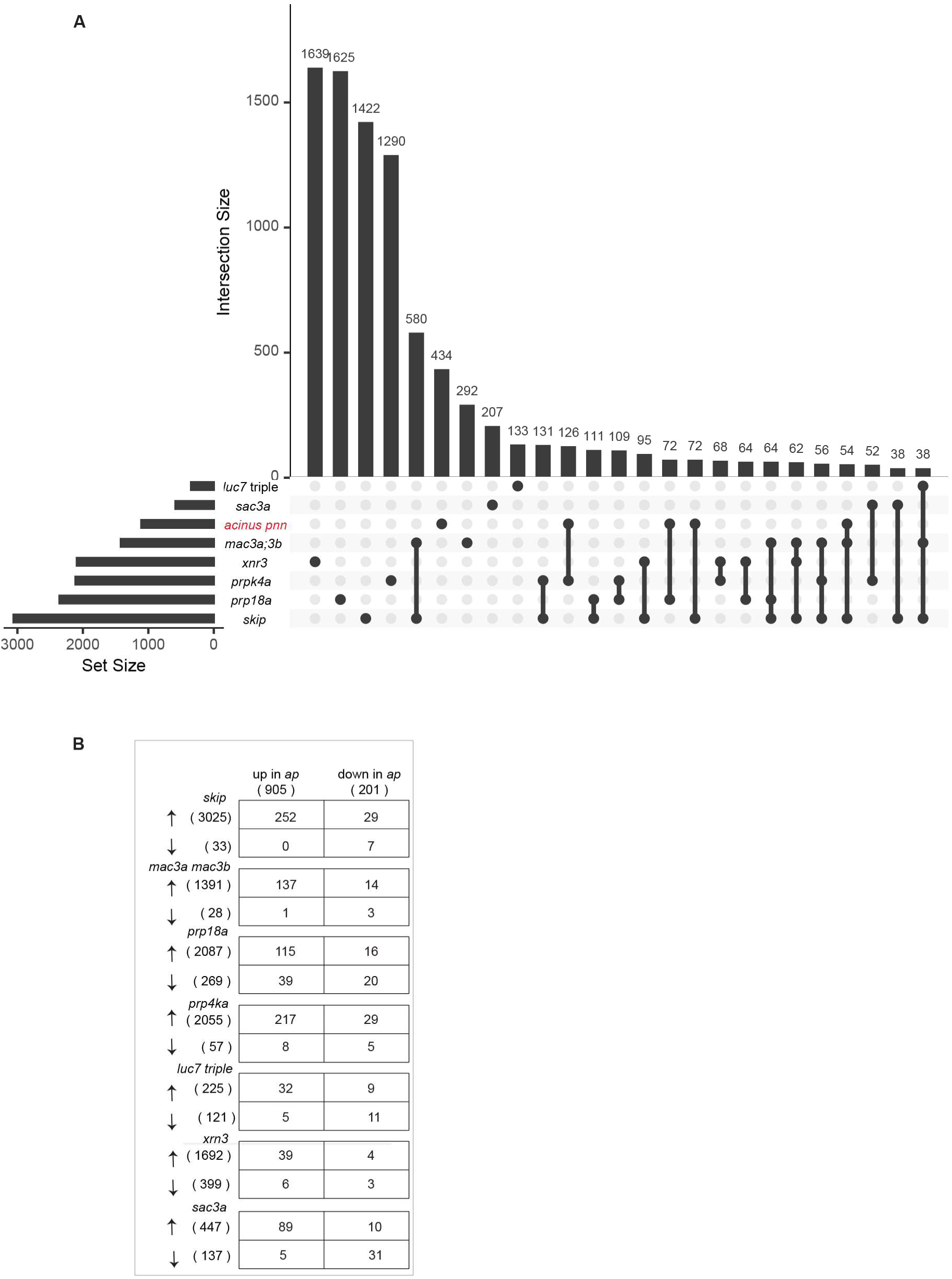
Common and distinct retained intron events regulated by components in the ACINUS network. (A) UpSet plot shows the overlapping and distinct retained intron events across all selected mutants. (B) Directionality of regulated retained intron events, comparing *acinus pinin* mutants with other selected mutants.

**Supplemental Fig. S7.**
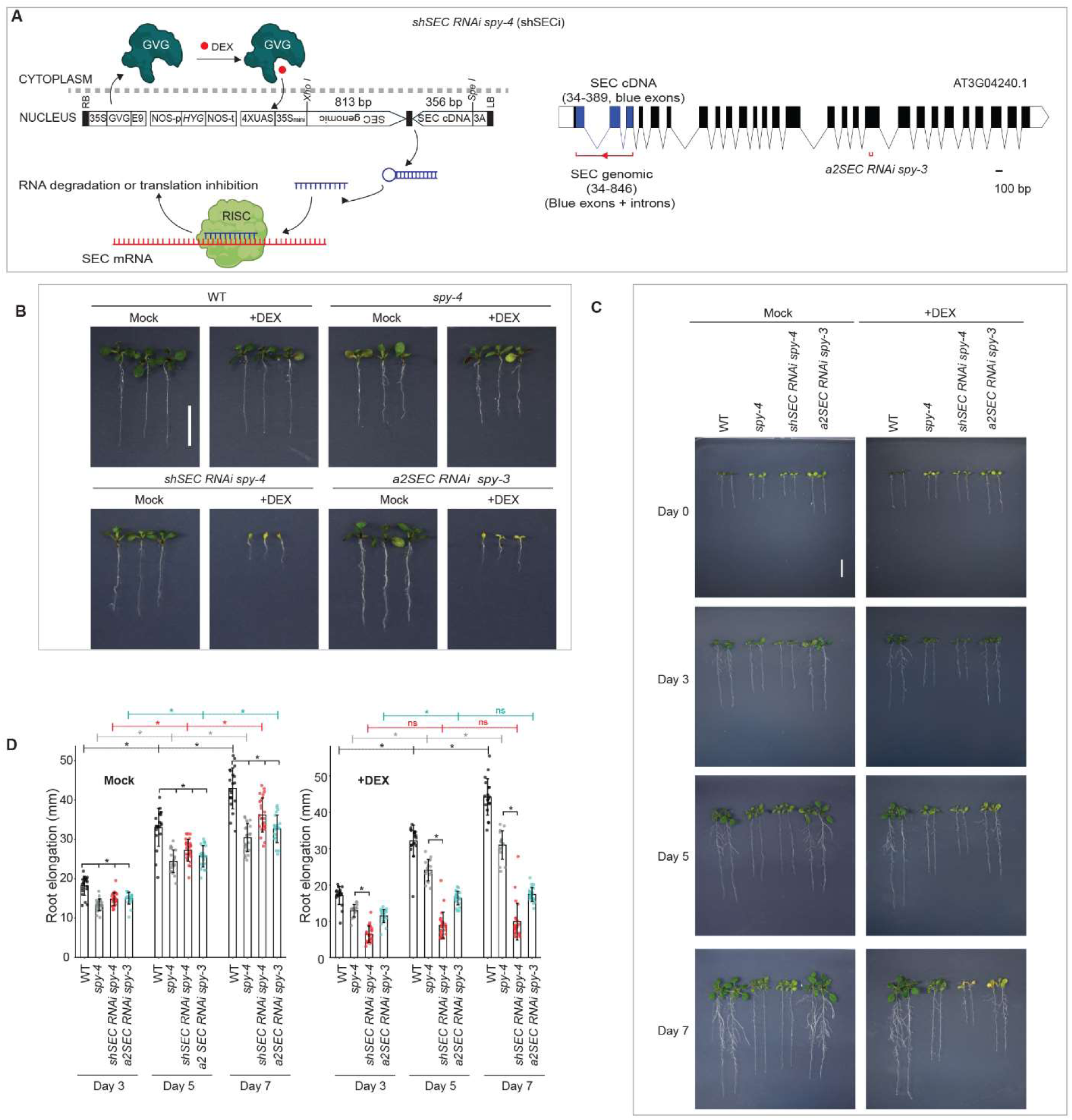
The inducible *shSEC RNAi spy-4* line shows seedling lethality. (A) Schematic of Dex-inducible *shSEC RNAi* construct compared to *a2SEC RNAi*. Genomic and cDNA of SEC segments are cloned in a head-to-head orientation to pTA7002 to generate the short hairpin RNAi. (B) Seedling lethality was observed when double mutants were grown directly on Dex-supplemented plates. (C) Progressive phenotype of *shSEC RNAi spy-4* phenotype after Dex treatment, showing reduced root growth by day 3, leaf discoloration by day 5 and complete growth arrest by day 7. (**D**) Quantification of root elongation after transfer on mock and Dex treatment. Root lengths were measured in over 15 seedlings, and statistical analysis was performed using a two-tailed t-test (p < 0.001).

**Supplementary Fig. S8.**
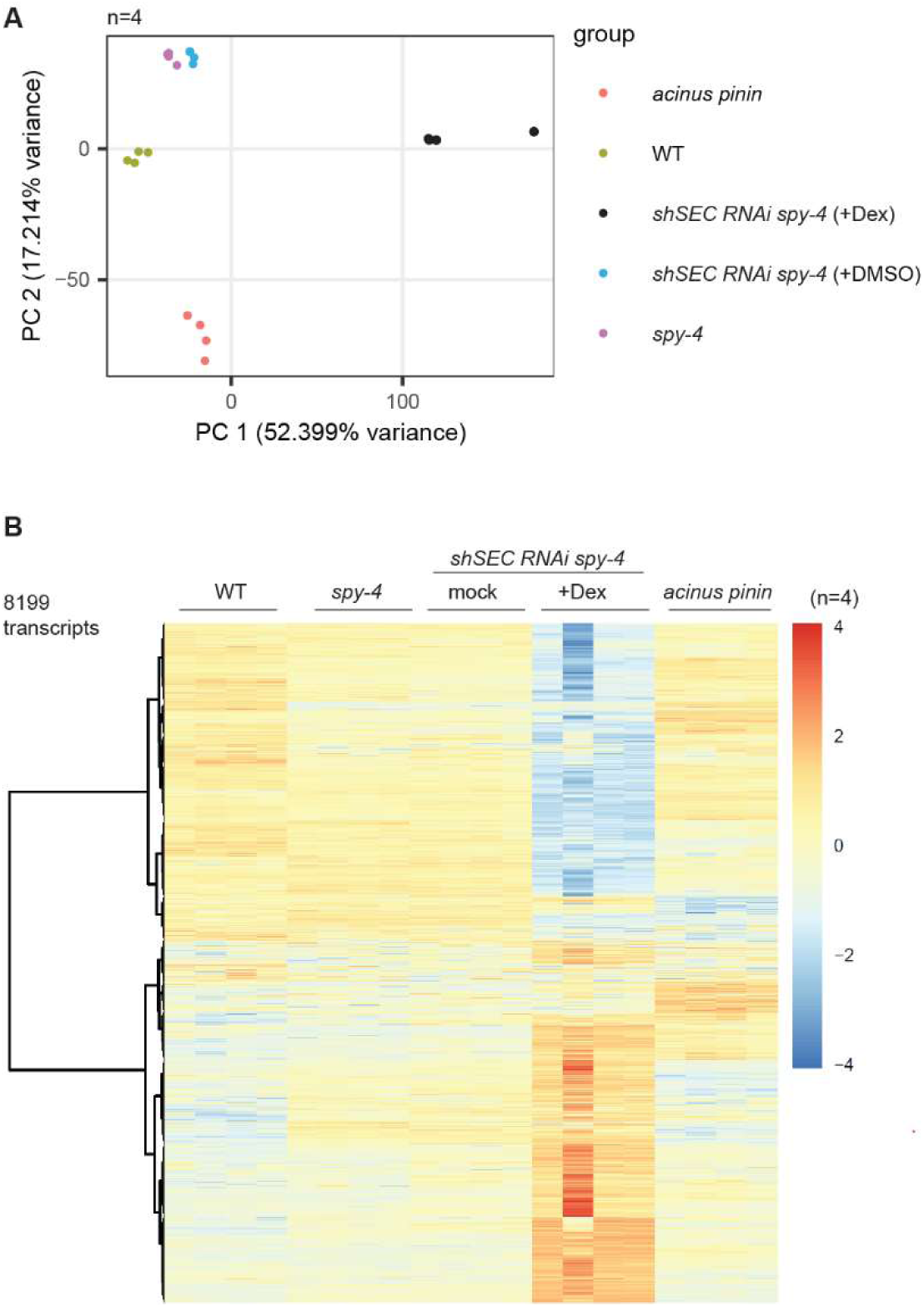
PCA analysis of RNA-seq data shows high reproducibility of data sets. (A) PCA clustering shows that the replicates of the WT, *acinus pinin, spy-4,* and *shSEC RNAi* groups clustered closely together, indicating high reproducibility, with minimal effects of DMSO treatment on seedlings. (B) Heatmap showing 8,199 differentially expressed transcripts from Fig. 7D that were altered in the *acinus pinin* mutant or *shSEC RNAi spy-4* with Dex treatment, or both conditions compared to WT. A total of 26,186 transcripts were quantified.

## Supplementary Table

**Supplementary Table S1: RNA-seq analysis result. Supplementary Table S2: Primers used in this study.**

**Supplementary Table S2: TurboID-related results.** The table includes Table content TurboID summary, evolution conserveness, ACINUS-TD_Protein-level result, PNN-TD_Protein-level result, SR45-TD_Protein-level result, Comparison of ACINUS-TD vs ACINUSΔRSB-TD, ACINUS-TD_Peptide Level, Interpro Results, and Interpro results counts.

**Supplementary Table S3: Primers used in this study.**

## Notes

### Competing Interest Statement

The authors have declared no competing interest.

### Summary of Updates

Two errors in the numbers listed in Table 1 have been corrected.

